# Micropolarity governs the structural organization of biomolecular condensates

**DOI:** 10.1101/2023.03.30.534881

**Authors:** Songtao Ye, Andrew P. Latham, Yuqi Tang, Chia-Heng Hsiung, Junlin Chen, Feng Luo, Yu Liu, Bin Zhang, Xin Zhang

## Abstract

Microenvironment is critical to the function of cells and organisms. One example is provided by biomolecular condensates, whose microenvironment can be vastly different from the surrounding cellular environments to engage unique biological functions. How microenvironments of biomolecular condensates affect their structure and function remains unknown. Here, we show that the arrangements and partitioning of biomolecules are dictated by the differences between the micropolarity of each subcompartment. Sufficient difference in micropolarity results in layered structures with the exterior shell presenting a more polar microenvironment than the interior core. Accordingly, micropolarity inversion is accompanied by conversions of the layered structures. These findings demonstrated the central role of the previously overlooked microenvironment in regulating the structural organization and function of membraneless organelles.

## INTRODUCTION

Biomolecular condensates are membraneless intracellular protein- and RNA-rich compartments that originated through a phase transition process termed liquid-liquid phase separation (LLPS) (*1*–*5*). The microenvironment inside the biomolecular condensates is drastically different from the outside surroundings, with specific properties including pH (*6*), biomacromolecule compactness (*7*), hydration (*8*, *9*), viscoelasticity (*10*, *11*), drug recruitment (*12*) and others. Till now, it remains elusive how microenvironments affect the structure and client recruitment of biomolecular condensates to allow their cellular function. This question becomes particularly intriguing for multiphasic biomolecular condensates that arrange in a “phase within phase” multiphasic structure, represented by stress granules (*13*, *14*), nucleoli (*15*–*17*), and anisosomes (*18*), amongst many others. While interfacial tension has been proposed to determine the arrangements of the layered structures (*16*, *17*, *19*), chemical principles to explain the regulation of multiphasic membraneless organelles calls remain less understood. For instance, why do biomolecules partition in different layers and what factor determines their partition? Answers to these questions would help understand the biological functions of the multiphasic membraneless organelles, as they recruit different ensembles of biomacromolecules and substrates to each layer.

Herein, we establish the innate biophysical property in the microenvironment of each subcompartment that controls the dynamics of structural organization and substrate partitioning of multiphasic membraneless organelles. As an *in vitro* model system, protein-rich condensates formed by elastin-like polypeptides (ELPs) were applied to investigate their microenvironments and how the microenvironment affects the organization and partitioning of multiphasic protein-rich condensates. The coacervation of two ELPs resulted in the formation of dual-component ELP condensates with diverse structures and partitioning results. We found that the micropolarity of the droplets’ microenvironment governs the structural arrangements of multiphasic ELP condensates. Tuning the organization of condensates was achieved through perturbing their microenvironments. Using nucleolus as a model multiphasic membraneless organelle, we found that the outermost granular component (GC) layer showed a more polar microenvironment than the inner dense fibrillar component (DFC) layer. Inhibition of RNA synthesis led to the dislocation of DFC to reside outside of GC layer, forming nucleolar caps. The inversion of polarity ranking between GC and DFC was found to be accompanied by the DFC dislocation. Collectively, these results demonstrated that micropolarity plays a central role in the organization and substrate partitioning of multiphasic protein-rich droplets and membraneless organelles. Spatiotemporal modulation of the multiphasic membraneless organelles in live cells is potentially achieved by tuning their microenvironment upon specific cell signaling, enabled by the influx and outflux of biomacromolecules and small molecule substrates.

## RESULTS

### Binary mixtures of ELP coacervate into condensates with diverse structures and miscibility

ELPs are bioengineered low-complexity peptide polymers that encode multiple pentameric repeats of VPGXG, in which the guest residue, X, could be any amino acid residue but proline (*20*). Previous work showed that the coacervation of multiple ELPs with different guest residues could form “phase within phase” condensates (*21*) (Fig. 1A). In this study, we generated purified ELPs bearing different guest residues and chain lengths (Fig. 1B). V_2_I_7_E-40 contains 40 pentameric repeats separated into 4 identical subgroups of 10 pentamers. For each subgroup, the pentamers bearing the guest residue of valine, isoleucine and glutamic acid were found at a ratio of 2:7:1. Likewise, KV_6_-112 and QV_6_-112 consist of 112 pentameric repeats that split evenly into 16 subgroups, with each subgroup made up of 1 pentamer bearing lysine or glutamine and 6 pentamers bearing valine. V-120, V_5_A_5_-120 and V_5_A_2_G_3_-120 are ELPs consisting of only aliphatic residues and glycine. V-120 is solely made up of pentamers bearing valine guest residue. V_5_A_5_-120 and V_5_A_2_G_3_-120 replaced half the number of valines with alanine and/or glycine residues.

**Figure 1.**
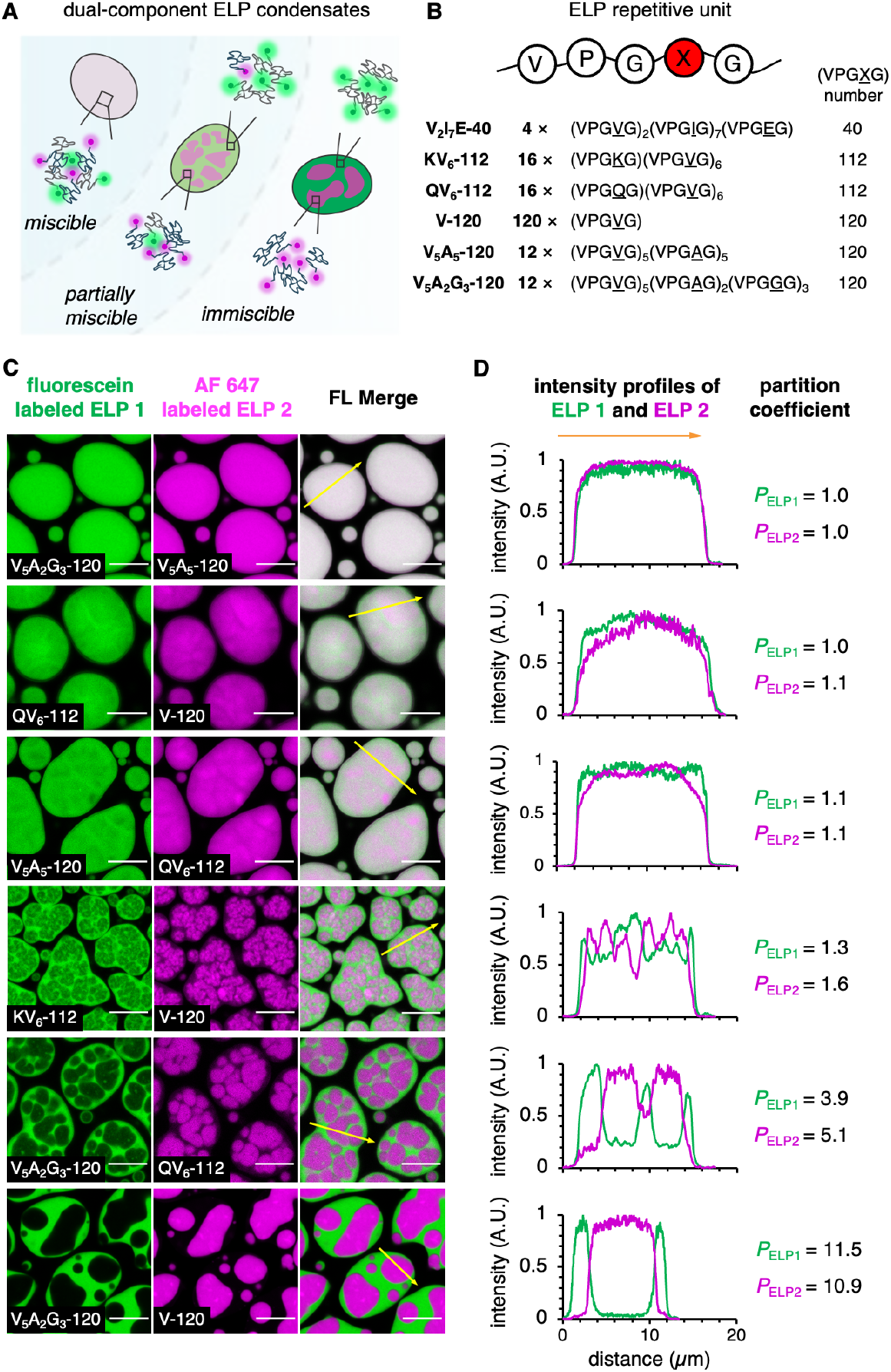
ELP binary mixtures phase separate into dual-component condensates with diverse phase behavior. **(A)** Schematic of the dual-component ELP condensates with diverse organization and miscibility. **(B)** The pentameric repetitive unit of ELP and the sequences of ELPs used in this work. **(C)** Dual-color confocal images of phase-separated dual-component ELP condensates. Co-phase separation of fluorescein-labeled ELP 1 (green) and AF 647-labeled ELP 2 (magenta) resulted in both miscible ELP condensates and dual-layer ELP droplets with various miscibility. Scale bars, 10 μm. **(D)** Fluorescence intensity profile along the yellow arrows from **C**. Partition coefficient values were calculated as the averaged fluorescence intensity ratio of ELPs between their rich and poor subcompartments in dual-component ELP droplets.

All purified ELPs display lower critical solution temperature phase behavior in aqueous solutions (*22*–*24*). ELPs undergo LLPS above phase transition temperatures (fig. S1), forming liquid droplets with merging and wetting properties (fig. S2). Next, we examined the coacervates of binary ELPs combinations by labeling the N-terminus of ELPs with either green-fluorescence fluorescein NHS-ester or red-fluorescence AF 647 NHS-ester. Interestingly, we observed different phase behaviors (Fig. 1C and fig. S3). Some binary combinations of ELPs appeared to be fully miscible with each other, represented by V_5_A_2_G_3_-120/V_5_A_5_-120. Others showed fluorescence heterogeneities, such as QV_6_-112/V-120 and V_5_A_5_-120/QV_6_-112 (Fig. 1C). Such heterogeneity became more pronounced in KV_6_-112/V-120 coacervates, where KV_6_-112 and V-120 were enriched in two subcompartments with clear boundaries. Finally, the V_5_A_2_G_3_-120/QV_6_-112 and V_5_A_2_G_3_-120/V-120 droplets presented predominantly separated core-shell structures, where each ELP remained in either the core or shell layer. Intensity profile analysis along the indicated lines from fluorescence images corroborated the findings of diverse phase behavior from dual-component ELP droplets (Fig. 1D). The partition coefficient (*P*_ELP_) of fluorescein-labeled ELP (indicated as ELP1) and AF 647 labeled ELP (ELP2) also increased from 1 for miscible V_5_A_2_G_3_-120/V_5_A_5_-120 droplets to over 10 for immiscible V_5_A_2_G_3_-120/V-120 droplets.

Several lines of evidence support the above observations were autonomous demixing of ELPs into a gradient of miscibility. First, in the LLPS process of dual-component ELP mixtures, as the solution temperature increases, ELP with lower T_ph_ initiates LLPS, followed by the demixing of another ELP with higher T_ph_ (movie S1). Secondly, the organization of the core-shell structure of ELPs was not affected by the sequence of phase separation, as the mixing of two already phase-separated ELPs resulted in similar structures as coacervated ELPs (fig. S4). Finally, swapping fluorophore labels on ELPs yields the same organization with inverted fluorescence colors (fig. S5), suggesting that the fluorophore labeling does not alter the organization nor the partitioning of the multi-component ELP condensates. These findings supported that the partially miscible and immiscible ELP condensates adopted a “phase within phase” architecture, resembling multiphasic membraneless organelles in cells.

### Micropolarity correlates to the organization and miscibility of the ELP condensates

Notably, we observed unexpected miscibility that contradicts previous understanding. For instance, the hydrophobic V-120 is fully miscible with negatively charged V_2_I_7_E-40, and partially miscible with positively charged KV_6_-112, yet surprisingly immiscible with non-charged V_5_A_2_G_3_-120. These results suggested that the miscibility of the dual-layered ELP condensates is not solely determined by the nature of their side chain hydrophobicity or charge. We then hypothesize that the microenvironment would be responsible for the formation of multiphasic condensates. To interrogate the biophysical properties of ELP droplets, we integrated the environmentally sensitive fluorophores with fluorescence lifetime imaging microscopy (FLIM) to quantify the microenvironment of condensates. The 7-sulfonamide benzoxadiazole fluorophore (hereinafter, SBD; Fig. 2A, left panel) was used to report micropolarity owing to its increasing fluorescence lifetime with decreasing polarity (*25*, *26*). A molecular rotor-based boron dipyrromethene (BODIPY) fluorophore was used to measure microviscosity (Fig. 2A, right panel), as its fluorescence lifetime elongates with increasing viscosity (*27*).

**Figure 2.**
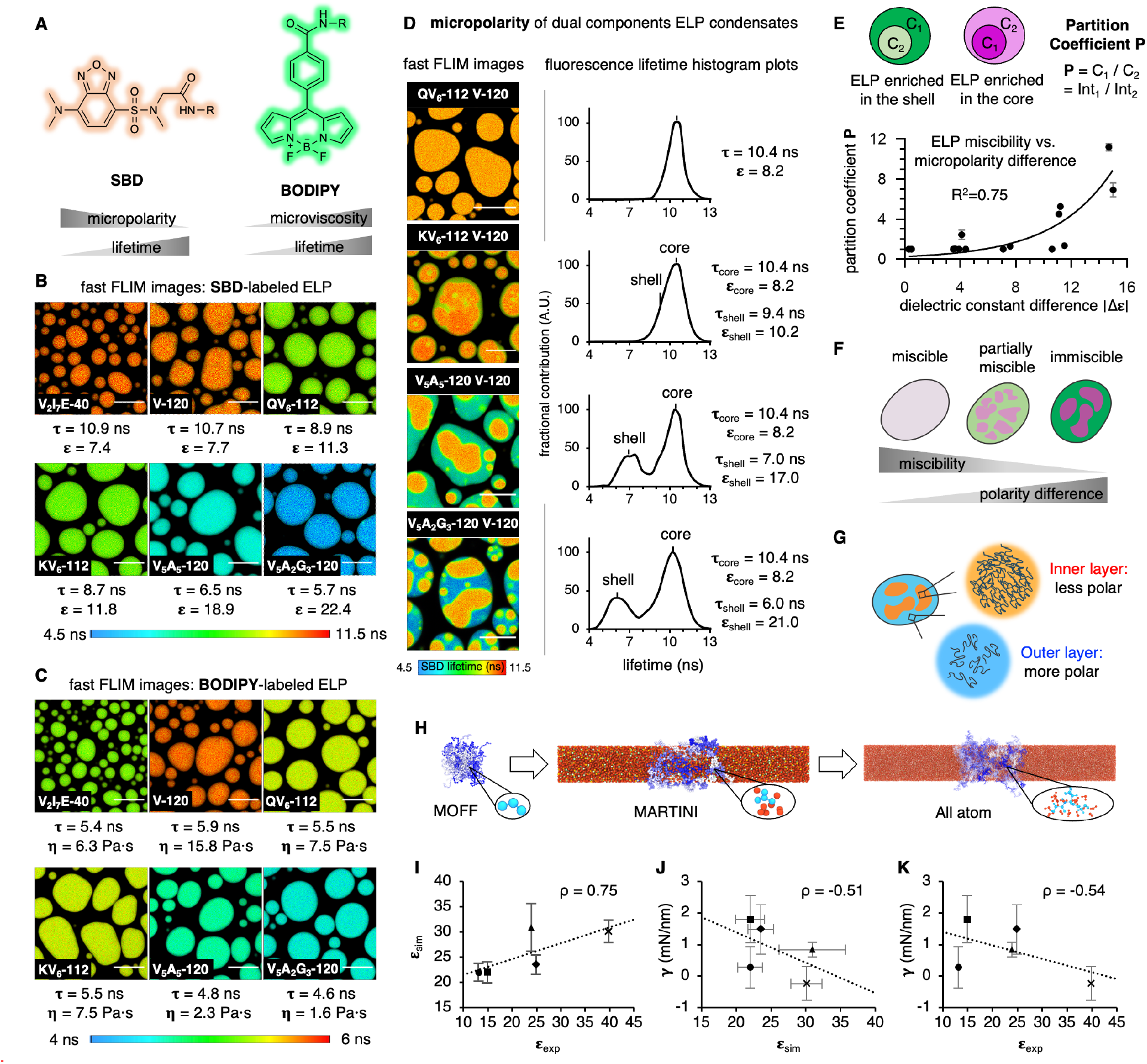
Polarity governs the organization and partition of the multi-component protein condensates. **(A)** Chemical structures of SBD and BODIPY. **(B-C)** FLIM images of SBD-**(B)** or BODIPY-**(C)** labeled ELP condensates. **(D)** FLIM images and histogram plots of dual SBD-labeled binary ELP condensates. Scale bars in **B-D**, 10 μm. **(E)** Relationship between the partition coefficient and the dielectric constant differences of binary ELP condensates. **(F)** Schematics showing the relationship between miscibility and polarity differences between two subcompartments of ELP condensates. **(G)** Schematics showing the polarity differences between the inner and the outer layer of multiphasic ELP droplets. **(H)** Multi-scale simulations used to simulate the surface tension and dielectric constant of ELP condensates. Surface tension is calculated using the MARTINI model (*35*), and dielectric constant is calculated using all-atom simulations (*36*). **(I)** Comparison between the experimental dielectric constant (ε_exp_) and the simulated dielectric constant (ε_sim_). **(J)** Comparison between ε_sim_ and the simulated surface tension (**γ**). **(K)** Comparison between the ε_exp_ and **γ**. In **I-K:** figure legends, dots, V_2_I_7_E^0^; squares, V_10_; triangles, QV_6_QV_2_; diamonds, K^0^V_6_K^0^V_2_; crosses, V_5_A_2_G_3_. The superscript 0 indicates non-protonated K and protonated E (Table S1). ρ, Pearson correlation coefficient. Error bars, standard diviations from 5 independent time windows (τ) or from 3 different starting configurations (ε).

We first measured the micropolarity and microviscosity of individual ELP droplets, using SBD or BODIPY, labeled on the N-terminus of ELP via amine-NHS chemistry. For SBD-labeled ELP droplets, the fluorescence lifetime spans from 5.7 ns for V_5_A_2_G_3_-120 to 10.9 ns for V_2_I_7_E-40 (Fig. 2B and figs. S6-8). Using the characteristic intensity-weighted mean fluorescence lifetime (τ_AvInt_), we converted the fluorescence lifetime of SBD to relating dielectric constant (ε) values (fig. S7) (*28*). The V_2_I_7_E-40 and V-120 droplets demonstrated surprisingly non-polar environments (ε = 7.4 and 7.7), despite that V_2_I_7_E-40 contained the charged residue glutamic acid. While no charged or polar residues were found within V_5_A_5_-120 and V_5_A_2_G_3_-120, they formed the most polar droplets with ε as 18.9 and 22.4, respectively. The microviscosity of the ELP constructs was also measured using BODIPY (Fig. 2C and fig. S9), whose fluorescence lifetime was correlated to viscosity values via a standard curve (fig. S10). V_5_A_2_G_3_-120 and V_5_A_5_-120 showed the least viscous microenvironment with the viscosity (η) calculated to be 1.6 Pa·s and 2.3 Pa·s, respectively. Whereas V-120 exhibited the most viscous environment, with its viscosity calculated to be 15.8 Pa·s. The observed viscosity values are consistent with the recovery rate from FRAP experiments (fig. S11).

We then focused on ELP coacervates and quantified the micropolarity and microviscosity of the outer shell and the inner core. When representative biphasic ELP condensates were prepared by mixing V-120 with QV_6_-112, KV_6_-112, V_5_A_5_-120 and V_5_A_2_G_3_-120 individually, we found a wide distribution of miscibility (Fig. 2D). The pseudo-color FLIM image of QV_6_-112/V-120 droplets demonstrated a uniform dielectric constant around 8.2 (Fig. 2D, top panel). By contrast, the KV_6_-112/V-120 droplets formed partially miscible layers (Fig. 2D, second panel), exhibiting a less polar core (ε = 8.2) surrounded by a polar shell (ε = 10.2). The V_5_A_5_-120/V-120 and V_5_A_2_G_3_-120/V-120 droplets further increased the micropolarity gap between the core and the shell (Fig. 2D, bottom 2 panels), wherein the dielectric constant values of the shell and the core were maximized to be 21.0 (shell) and 8.2 (core) for the V_5_A_2_G_3_-120/V-120 droplets. Furthermore, microviscosity measurements using the BODIPY probe also revealed a more viscous core than the shell (fig. S12), indicating that the microenvironment of the biphasic ELP droplets has a profound impact on their organization and partitioning.

We also evaluated whether polarity or viscosity is more important to the partitioning of biphasic ELP droplets. To this end, we calculated the partition coefficient (P) of the component ELPs from dual-color confocal images. The P values were plotted against the difference in polarity (Δε) or viscosity (Δη) of the individually formed ELP droplets under identical conditions and fitted with exponential curves. For polarity, we observed a positive correlation between the P values and Δε (Fig. 2E, R^2^ = 0.75), suggesting that the miscibility of the biphasic ELP condensates is negatively correlated with the polarity differences between the two components. By contrast, Δη exhibited a poor correlation with the P values (fig. S12, R^2^ = 0.52), suggesting the micropolarity governs the organization and partition of the multiphasic protein condensates (Fig. 2F). Furthermore, the shell layer harbors a more polar microenvironment than the core layer (Fig. 2G).

Interfacial tension was previously reported to play an important role in the structural organization of multiphasic condensates, albeit the correlation between interfacial tension and the physiochemical property remains elusive (*16*, *17*, *19*). To examine the interfacial tension of ELP condensates, we measured their contact angles on hydrophilic surfaces. Non-polar V-120 droplet exhibited greater interfacial tension, reflected by its larger contact angles of 122° ± 8° on hydrophilic glass surfaces, than polar ELP condensates (contact angles between 110° - 114°) (fig. S13). To explore this relationship further, we employed a multiscale simulation approach. Utilizing an all-atom representation, we calculated the dielectric constant (ε_sim_) of droplets formed by simplified 10 amino-acid ELP fragments (Fig. 2H and fig. S14), resulting in similar values compared with experimental dielectric constants (ε_exp_) that were acquired under the same condition with simulation (Fig. 2I and fig. S15; Table S1). Our simulations further suggest that the charged residues may adopt neutral states as a result of the more hydrophobic condensate interior, resulting in non-protonated K and protonated E. From our multiscale model, we also calculated the surface tension (γ_sim_) of ELP droplets (Table S1), revealing negative correlations between γ_sim_ and dielectric constant, either ε_sim_ (Fig. 2J) or ε_exp_ (Fig. 2K). Taken together, these results suggest that the micropolarity of ELP droplets dictates their interfacial tension, thus controlling the structural organization and partition of multiphasic ELP condensates.

### Micropolarity perturbation corresponds to structural changes of multiphasic condensates

Our results marked the importance of micropolarity in governing the structure and partitioning of multiphasic protein condensates. If this principle holds, then the perturbation of micropolarity should change the organization of the multiphasic condensates (Fig. 3A). To test this notion, we assembled coacervates of AF 647-labeled V_2_I_7_E-40 and fluorescein-labeled QV_6_-112, which formed fully miscible droplets with a homogenous micropolarity (τ = 3.9 ns and ε = 32.9; Fig. 3B-C, left panel). We used polyR, a 12-residue arginine repeat, to modulate the QV_6_-112/V_2_I_7_E-40 condensate, because polyR could interact with V_2_I_7_E-40 via electrostatic interaction and minimally interact with QV_6_-112 (fig. S16). When coumarin-labeled polyR was introduced to the V_2_I_7_E-40/QV_6_-112 droplet, we observed the miscible condensates segregate into punctuated V_2_I_7_E-40/polyR enriched cores and QV_6_-112 enriched shell (Fig. 3B, right panel). Subsequential FLIM imaging (Fig. 3C) and fluorescence lifetime histogram analysis (Fig. 3D) showed the emergence of less polar (longer SBD lifetime) V_2_I_7_E-40/polyR-enriched cores surrounded by QV_6_-112-enriched shell. These results demonstrated a “push-in” model, wherein micropolarity reduction of the newly formed phase could drive its segregation from miscible condensates, forming more hydrophobic cores inside the shells (Fig. 3A, “push-in” model).

**Figure 3.**
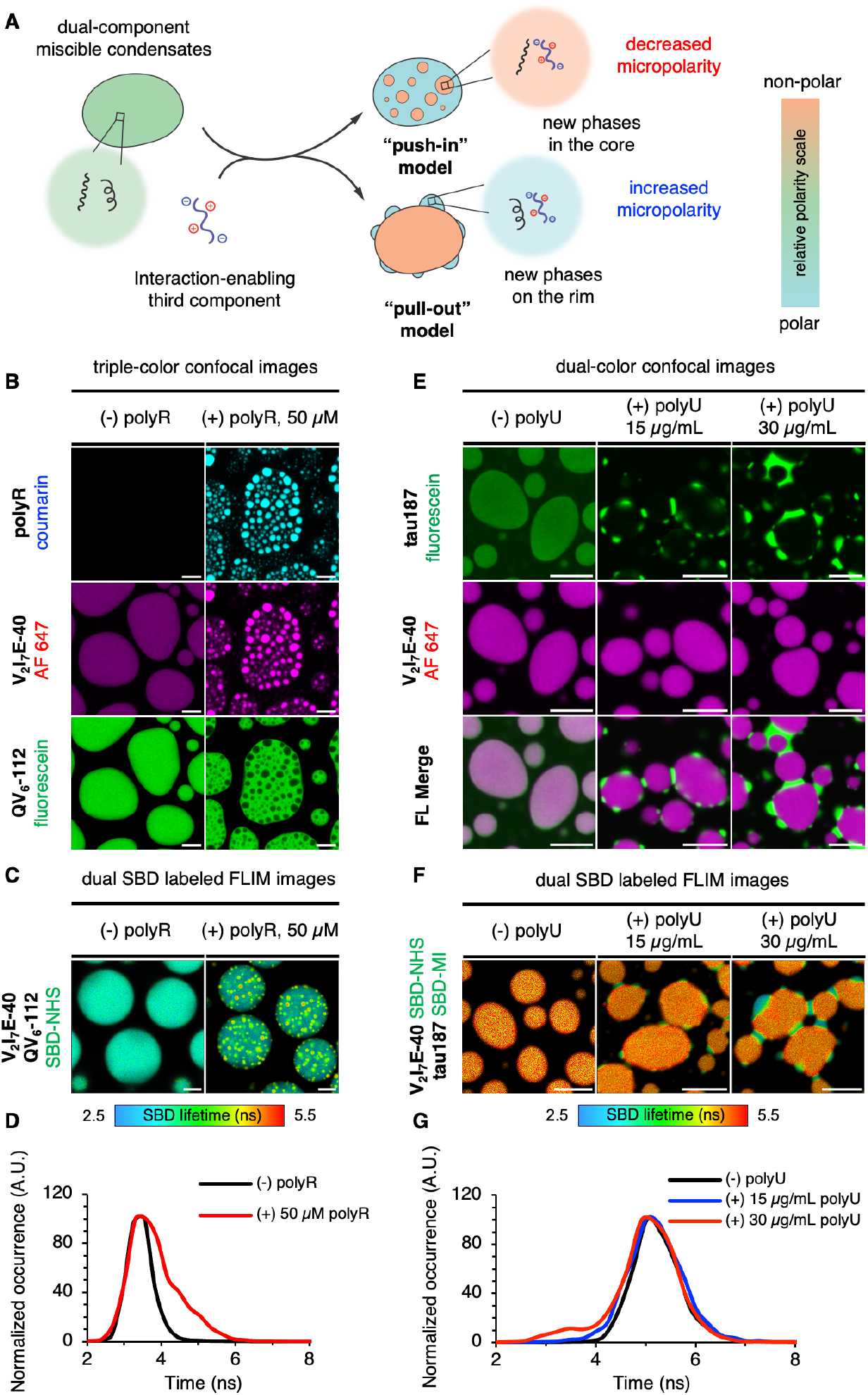
Modulating the structural organization of multi-component protein condensates through tuning micropolarity. **(A)** Illustration of the de-mixing of miscible protein condensates into multi-subcompartment protein condensates by the addition of a third interaction-enabling component. **(B)** Triple-color confocal images of protein condensates formed by AF 647-labeled V_2_I_7_E-40 and fluorescein-labeled QV_6_-112 in the absence or presence of coumarin-labeled polyR. **(C)** FLIM images of protein condensates formed by SBD-labeled V_2_I_7_E-40 and SBD-labeled QV_6_-112 in the absence or presence of polyR. **(D)** Fluorescence lifetime histogram plot for images in **C**. **(E)** Dual-color confocal images of protein condensates formed by fluorescein-labeled tau187 and AF 647-labeled V_2_I_7_E-40 in the absence or presence of different concentrations of polyU. **(F)** FLIM images of protein condensates formed by SBD-labeled tau187 and SBD-labeled V_2_I_7_E-40 in the absence or presence of different concentration of polyU. **(G)** Fluorescence lifetime histogram plot for images in **F**. Scale bars for images in **B-C** and **E-F,** 5 μm.

We next examined whether micropolarity elevation would lead to the formation of shells from miscible droplets. To this end, we generated coacervates of V_2_I_7_E-40 and the 187 amino-acid C-terminal half of the longest isoform of tau 2N4R proteins (hereinafter named tau187). The tau187/V_2_I_7_E-40 droplet was found to be fully miscible, with a homogeneous micropolarity and a dielectric constant around 24.9 (τ = 5.2 ns, Fig. 3E-F, left panel). It has been reported that tau187 interacts with RNA to form tau/RNA coacervates (fig. S17A-E) (*29*, *30*). When we added 15 μg/mL polyU to the tau187/V_2_I_7_E-40 droplet, tau187 formed coacervation with polyU and was enriched on the outer rim of the V_2_I_7_E-40 condensates (Fig. 3E, middle panel). Higher polyU concentration led to the expansion of the outer tau187 enriched layer to a greater extent, forming condensed puncta attached to the outer periphery of V_2_I_7_E-40-enriched condensates (Fig. 3E, right panel). Accordingly, FLIM experiments revealed increasing micropolarity of the tau187 enriched puncta with increasing concentrations of polyU (fig. S17F), with the dielectric constant of tau187 changing from 24.9 to 35.8 (τ = 3.5 ns; Figs. 3F-G). Thus, adding polyU increased the polarity of tau187 and pulled it out from the tau187/V_2_I_7_E-40 droplet to the tau187/polyU shell, leaving the non-polar V_2_I_7_E-40 droplet as the core structure (Fig. 3A, “pull-out” model).

### Micropolarity controls the structural features of the multiphasic nucleolus

To verify whether micropolarity could dictate the structure of the cellular multiphasic membraneless organelles, we investigated the microenvironment of the multilayered nucleolus in eukaryotic cells. Nucleolus consists of three distinct phases: fibrillar center (FC), dense fibrillar component (DFC), and granular component (GC) (*15*–*17*, *31*). Structurally, the FC is embedded inside the DFC, and the FC-DFC assembly is further encapsulated by the GC layer (Fig. 4A-B). First, we confirmed the hierarchical structure of nucleolus in HEK293T cells by co-expression of the GC component nucleophosmin (EGFP-NPM1) and the DFC component fibrillarin (FBL-Halo), labeled with the HaloTag^®^ TMR ligand (Fig. 4A, left panel). When transcription was inhibited by the Pol I inhibitor, actinomycin D (ActD), the FC and DFC layers were found to be spatiotemporally translocated to the outside and juxtaposed on the peripheral of the GC layer, forming nucleolar caps (Fig. 4A-B) (*32*, *33*).

**Figure 4.**
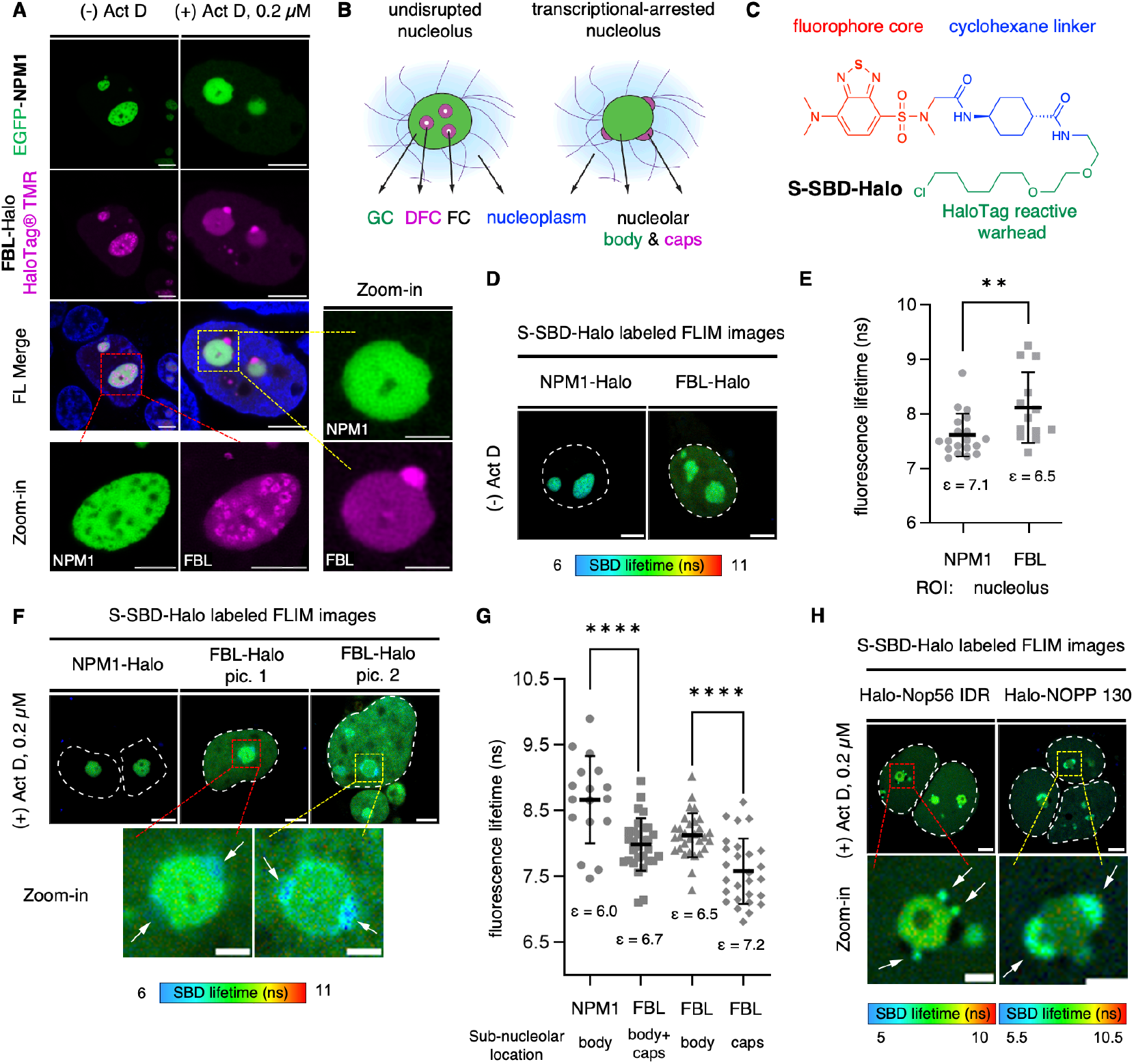
Micropolarity measurements in healthy and transcriptional-arrested nucleolus revealed a micropolarity inversion between DFC and GC subcompartments accompanied with organization change. **(A)** Triple-color imaging of HEK293T cells co-expressing EGFP-NPM1 and FBL-Halo in the presence or absence of actinomycin D (Act D). **(B)** Multiphasic structures of the nucleolar subcompartments. **(C)** Chemical structure of S-SBD-Halo fluorophore. **(D-E)** Representative FLIM images and fluorescence lifetime (τ_AvInt_) statistics of cells expressing NPMl-Halo or FBL-Halo labeled by S-SBD-Halo in the absence of Act D. n = 19 for NPM1 and n = 13 for FBL in **E**. **(F-G)** Representative FLIM images and τ_AvInt_ statistics of HEK 293T cells expressing NPM1-Halo or FBL-Halo labeled by S-SBD-Halo probes in the presence of Act D. n = 17 for NPM1, n = 30 for FBL in the entire nucleolus, n = 30 for FBL in the nucleolar body and n = 27 for FBL in the nucleolar caps in **G**. **(H)** FLIM images of HEK 293T cells expressing Halo-Nop56 IDR or Halo-NOPP 130 labeled by S-SBD-Halo probes in the presence of 0.2 μM Act D. Scale bars, 2 μm for zoom-in figures and 5 μm for others. Statistical significance was calculated using two tailed t-test. **0.01<P<0.001, ****P<0.0001.

To capture the nucleolus micropolarity changes associated with this structural inversion, we synthesized the SBD-Halo ligand (S-SBD-Halo) to enable the live-cell FLIM experiment (Fig. 4C and fig. S18). Firstly, we measured the micropolarity value surrounding NPM1 and FBL as representations of the overall micropolarity profiles for GC and DFC layers. To this end, we expressed NPM1-Halo or FBL-Halo in HEK293T cells in the presence of S-SBD-Halo to determine the micropolarity of the GC and DFC regions (Fig. 4D-E). The dielectric constant values of the GC and DFC regions were determined to be 7.1 (τ = 7.6 ns; n = 19) and 6.5 (τ = 8.1 ns; n = 13), respectively (Fig. 4E). These data reflect that the GC layer harbors a slightly more polar microenvironment compared to the DFC layer. In the presence of Act D, the dielectric constant values of GC and DFC regions were 6.0 (τ = 8.7 ns; n = 17) and 6.7 (τ = 8.0 ns; n = 30), respectively (Figs. 4F-G), suggesting a micropolarity inversion of the GC and DFC layers. Importantly, FBL located in the nucleolar caps demonstrated a more polar microenvironment (ε = 7.2, τ = 7.6 ns; n = 27) than the FBL located in the nucleolar body (ε = 6.5, τ = 8.1 ns; n = 30), as determined by both representative FLIM images (Fig. 4F, zoom-in) and statistical analysis (Fig. 4G). In addition to NPM1 and FBL, we observed a similar trend using the DFC-enriched NOP56 protein (C-terminal intrinsically-disordered region, fused with HaloTag) (*34*), as well as the nucleolar phosphoprotein p130 (NOPP 130, encoded by NOLC1 gene). In both cases, the nucleolar cap regions demonstrated a shorter lifetime (hence a more polar microenvironment) compared to the rest of the nucleolar bodies (Fig. 4H). These findings provide direct evidence to demonstrate that the phase inversion of the nucleolus is directly associated with the micropolarity inversion between the GC and DFC layers.

## DISCUSSION

We have uncovered a chemical principle that the micropolarity governs the layered structures and substance partition of multiphasic condensates. We determined that the micropolarity negatively corresponds to the surface tension. And thus, the extent of micropolarity differences correlates to the extent of layered structures. Substances with a greater micropolarity partition into the exterior shell, and the interior core presents a more non-polar microenvironment. Accordingly, micropolarity inversion is accompanied by the conversion of the core-shell structures.

These results demonstrated the delicate control of the organization and partition of the multiphasic condensates by tuning the polarity of individual components. The dynamic interplay between the micropolarity and layered structures offers a new insight regarding multiphase organelle regulation in live cells. The multi-valent interaction between droplets’ component proteins with other proteins/RNAs influx into the droplets upon specific signals could change the overall micropolarity of the newly formed complexes, which destabilize the droplets by elevating the free energy of mixing and cause the organization and partitioning change. Our discovery provides a novel perspective regarding how multi-component biomolecular condensates are organized *in vitro* and in live cells. The understanding of condensates’ micropolarity, in combination with other physical properties such as microviscosity and recently proposed capillary forces, could collectively help us to probe the complex biophysical states of membraneless organelles, and ultimately deepen our understanding of their regulation and cellular functions.

## ACKNOWLEDGMENTS

The authors thank Instrumentation and Service Center for Molecular Sciences at Westlake University for the assistance on fluorescence lifetime imaging microscopy (FLIM). We thank the Research Center for Industries of the Future (RCIF) at Westlake University for supporting this work, which was supported by the National Natural Science Foundation of China (22222410). B.Z. and A.L. were supported by the National Institutes of Health (Grant R35GM133580). A.L. acknowledges support from the National Science Foundation Graduate Research Fellowship Program (grant number 1745302).

## AUTHOR CONTRIBUTIONS

S.Y. and X.Z. conceived the project; S.Y., Y.T., C.H., performed experiments; A.P.L performed computational analysis; J.C. assisted with plasmids cloning; F.L. assisted with organic synthesis and characterization; Y.L., B.Z., and X.Z., supervised the project. S.Y. and X.Z. wrote the paper with help from other coauthors.

## COMPETING INTERESTS

Authors declare that they have no competing interests.

## DATA AND MATERIALS AVAILABILITY

All data are available in the main text or the supplementary materials.

## Supplementary Materials

Materials and Methods

Figs. S1 to S18

Tables S1 to S4

Scheme S1

References (*36*-54)

Movies S1

## Notes

### Competing Interest Statement

The authors have declared no competing interest.

## References and Notes

1. C. P. Brangwynne, C. R. Eckmann, D. S. Courson, A. Rybarska, C. Hoege, J. Gharakhani, F. Jülicher, A. A. Hyman, Germline P granules are liquid droplets that localize by controlled dissolution/condensation. Science 324, 1729–1732 (2009).

2. A. A. Hyman, C. A. Weber, F. Jülicher, Liquid-liquid phase separation in biology. Annual Review Cell Developmental Biology 30, 39–58 (2014).

3. E. Gomes, J. Shorter, The molecular language of membraneless organelles. Journal of Biological Chemistry 294, 7115–7127 (2019).

4. S. F. Banani, H. O. Lee, A. A. Hyman, M. K. Rosen, Biomolecular condensates: organizers of cellular biochemistry. Nature Reviews Molecular Cell biology 18, 285–298 (2017).

5. Y. Shin, C. P. Brangwynne, Liquid phase condensation in cell physiology and disease. Science 357, eaaf4382 (2017).

6. P. Dogra, A. Joshi, A. Majumdar, S. Mukhopadhyay, Intermolecular charge-transfer modulates liquid–liquid phase separation and liquid-to-solid maturation of an intrinsically disordered pH-responsive domain. Journal of the American Chemical Society 141, 20380–20389 (2019).

7. A. Abyzov, M. Blackledge, M. Zweckstetter, Conformational dynamics of intrinsically disordered proteins regulate biomolecular condensate chemistry. Chemical Reviews 122, 6719–6748 (2022).

8. J. Ahlers, E. M. Adams, V. Bader, S. Pezzotti, K. F. Winklhofer, J. Tatzelt, M. Havenith, The key role of solvent in condensation: Mapping water in liquid-liquid phase-separated FUS. Biophysical Journal 120, 1266–1275 (2021).

9. A. P. Latham, B. Zhang, Molecular determinants for the layering and coarsening of biological condensates. Aggregate 3, e306 (2022).

10. L. Jawerth, E. Fischer-Friedrich, S. Saha, J. Wang, T. Franzmann, X. Zhang, J. Sachweh, M. Ruer, M. Ijavi, S. Saha, J. Mahamid, A. A. Hyman, F. Jülicher, Protein condensates as aging Maxwell fluids. Science 370, 1317–1323 (2020).

11. I. Alshareedah, M. M. Moosa, M. Pham, D. A. Potoyan, P. R. Banerjee, Programmable viscoelasticity in protein-RNA condensates with disordered sticker-spacer polypeptides. Nature Communications 12, 6620 (2021).

12. I. A. Klein, A. Boija, L. K. Afeyan, S. W. Hawken, M. Fan, A. Dall’Agnese, O. Oksuz, J. E. Henninger, K. Shrinivas, B. R. Sabari, I. Sagi, V. E. Clark, J. M. Platt, M. Kar, P. M. Mccall, A. V. Zamudio, J. C. Manteiga, E. L. Coffey, C. H. Li, N. M. Hannett, Y. E. Guo, T. M. Decker, T. I. Lee, T. Zhang, J. K. Weng, D. J. Taatjes, A. Chakraborty, P. A. Sharp. Y. T. Chang, A. A. Hyman, N. S. Gray, R. A. Young, Partitioning of cancer therapeutics in nuclear condensates. Science 368, 1386–1392 (2020).

13. S. Jain, J. R. Wheeler, R. W. Walters, A. Agrawal, A. Barsic, R. Parker, ATPase-modulated stress granules contain a diverse proteome and substructure. Cell 164, 487–498 (2016).

14. D. S. Protter, R. Parker, Principles and properties of stress granules. Trends in Cell Biology 26, 668–679 (2016).

15. F.M. Boisvert, S. van Koningsbruggen, J. Navascués, A. I. Lamond, The multifunctional nucleolus. Nature Reviews Molecular Cell Biology 8, 574–585 (2007).

16. M. Feric, N. Vaidya, T. S. Harmon, D. M. Mitrea, L. Zhu, T. M. Richardson, R. W. Kriwacki, R. V. Pappu, C. P. Brangwynne, Coexisting liquid phases underlie nucleolar subcompartments. Cell 165, 1686–1697 (2016).

17. D. L. Lafontaine, J. A. Riback, R. Bascetin, C. P. Brangwynne, The nucleolus as a multiphase liquid condensate. Nature Reviews Molecular Cell Biology 22, 165–182 (2021).

18. H. Yu, S. Lu, K. Gasior, D. Singh, S. Vazquez-Sanchez, O. Tapia, D. Toprani, M. S. Beccari, J. R. Yates III, S. Da Cruz, HSP70 chaperones RNA-free TDP-43 into anisotropic intranuclear liquid spherical shells. Science 371, eabb4309 (2021).

19. B. Gouveia, Y. Kim, J. W. Shaevitz, S. Petry, H. A. Stone, C. P. Brangwynne, Capillary forces generated by biomolecular condensates. Nature 609, 255–264 (2022).

20. S. R. MacEwan, A. Chilkoti, Elastin-like polypeptides: biomedical applications of tunable biopolymers. Peptide Science 94, 60–77 (2010).

21. J. R. Simon, N. J. Carroll, M. Rubinstein, A. Chilkoti, G. P. López, Programming molecular self-assembly of intrinsically disordered proteins containing sequences of low complexity. Nature Chemistry 9, 509–515 (2017).

22. Y. H. Cho, Y. J. Zhang, T. Christensen, L. B. Sagle, A. Chilkoti, P. S. Cremer, Effects of hofmeister anions on the phase transition temperature of elastin-like polypeptides. Journal of Physical Chemistry B 112, 13765–13771 (2008).

23. N. K. Li, F. G. Quiroz, C. K. Hall, A. Chilkoti, Y. G. Yingling, Molecular description of the LCST behavior of an elastin-like polypeptide. Biomacromolecules 15, 3522–3530 (2014).

24. D. W. Urry, Journal of Physical Chemistry B 101, 11007–11028 (1997).

25. K. H. Jung, S. F. Kim, Y. Liu, X. Zhang, A fluorogenic AggTag method based on Halo- and SNAP-tags to simultaneously detect aggregation of two proteins in live cells. ChemBioChem 20, 1078–1087 (2019).

26. Y. Liu, K. Miao, N. P. Dunham, H. Liu, M. Fares, A. K. Boal, X. Li, X. Zhang, The cation-π interaction enables a halo-tag fluorogenic probe for fast no-wash live cell imaging and gel-free protein quantification. Biochemistry 56, 1585–1595 (2017).

27. B. Shen, K. H. Jung, S. Ye, C. A. Hoelzel, C. H. Wolstenholme, H. Huang, Y. Liu, X. Zhang, A dual-functional BODIPY-based molecular rotor probe reveals different viscosity of protein aggregates in live cells. Aggregate DOI: 10.1002/agt2.301 (2022).

28. E. Fišerová, M. Kubala, Mean fluorescence lifetime and its error. Journal of Luminescence 132, 2059–2064 (2012).

29. Y. Lin, Y. Fichou, A. P. Longhini, L. C. Llanes, P. Yin, G. C. Bazan, K. S. Kosik, S. Han, Liquid-liquid phase separation of tau driven by hydrophobic interaction facilitates fibrillization of tau. Journal of Molecular Biology 433, 166731 (2021).

30. Y. Lin, J. McCarty, J. N. Rauch, K. T. Delaney, K. S. Kosik, G. H. Fredrickson, J.E. Shea, S. Han, Narrow equilibrium window for complex coacervation of tau and RNA under cellular conditions. Elife 8, e42571 (2019).

31. H. G. Schwarzacher, F. Wachtler, The nucleolus. Anatomy and embryology 188, 515–536 (1993).

32. R. Reynolds, P. O’B. Montgomery, B. Hughes, Nucleolar “caps” produced by actinomycin D. Cancer Research 24, 1269–1277 (1964).

33. Y. Shav-Tal, J. Blechman, X. Darzacq, C. Montagna, B. T. Dye, J. G. Patton, R. H. Singer, D. Zipori, Dynamic sorting of nuclear components into distinct nucleolar caps during transcriptional inhibition. Molecular Biology of the Cell 16, 2395–2413 (2005).

34. T. Gautier, T. Bergès, D. Tollervey, E. Hurt, Nucleolar KKE/D repeat proteins Nop56p and Nop58p interact with Nop1p and are required for ribosome biogenesis. Molecular and Cellular Biology 17, 7088–7098 (1997).

35. P. C. Souza, R. Alessandri, J. Barnoud, S. Thallmair, I. Faustino, F. Grünewald, I. Patmanidis, H. Abdizadeh, B. M. Bruininks, T. A. Wassenaar, P. C. Kroon, J. Melcr, V. Nieto, V. Corradi, H. M. Khan, J. Domański, M. Javanainen, H. Martinez-Seara, N. Reuter, R. B. Best, I. Vattulainen, L. Monticelli, X, Periole, D. P. Tieleman, A. H. de Vries, S. J. Marrink. Martini 3: a general purpose force field for coarse-grained molecular dynamics. Nature Methods 18, 382–388 (2021).

36. J. Huang, S. Rauscher, G. Nawrocki, T. Ran, M. Feig, B. L. De Groot, H. Grubmüller, A. D. MacKerell Jr., CHARMM36m: an improved force field for folded and intrinsically disordered proteins. Nature Methods 14, 71–73 (2017).

