## Supplementary Materials for "Micropolarity governs the structural organization of biomolecular condensates"

**This PDF file includes:**

Materials and Methods
Figs. S1 to S18
Tables S1 to S4
Scheme S1
References (*36–54*)
Characterization
Captions for Movies S1

**Other Supplementary Materials for this manuscript include the following:**

Movies S1

**Materials and Methods**

1. Construction of plasmids.

Construction of ELP plasmids:

The desired ELP sequences were constructed in pET-28a (+) vector using the previously reported Recursive Directional Ligation by Plasmids Reconstruction (Pre-RDL) method (*36*). All restriction enzymes were purchased from New England Biolabs® unless otherwise specified. The original pET-28a(+) vector (Novagen®) was first modified to insert BseRI and AcuI enzyme recognition sequences to allow Pre-RDL cloning. 1.5 μg pET-28a(+) plasmids were digested with XbaI and BamHI at 37˚C for 4 hrs, followed by dephosphorylation with rSAP at 37˚C for 2 hrs. The reaction mixture was purified with a PCR clean-up kit (Omega E.Z.N.A.® Cycle Pure Kit) to acquire the linearized vector. Two 5'-phosphorylated DNA oligos (oligo 1 and oligo 2, table S2) were synthesized and annealed at 2 μM concentration in T4 DNA ligation buffer by heating the reaction to 95˚C for 3 minutes, followed by stepwise cooling down at 0.25˚C/min rate to room temperature to resolve the insertion part. The vector and the insertion were mixed at a 1:3 molar ratio and incubated with T4 DNA ligase at room temperature for 2 hrs. The ligation product was transformed into Stbl3 chemically competent cells (Invitrogen$^{TM}$, C737303), allowed to recover for 1 hr in SOC at 37˚C. The cells were plated on LB agar plated supplied with 50 μg/mL of kanamycin sulfate. Single *E. coli* colony was picked, amplified and sequenced to resolve the correct plasmid, hereinafter named the modified pET 28a(+).

Two 75 bp 5'-phosphoralated DNA oligos encoding the ELP sequence (GXGVP-5, oligo 3 to oligo 20, table S2) were synthesized and annealed at 2 μM concentration in T4 DNA ligation buffer by heating the reaction to 95˚C and cooling slowly as described above. 1.5 μg of modified pET 28a(+) plasmid was digested with BseRI at 37˚C for 4 hrs, followed by dephosphorylation with rSAP at 37˚C for 2 hrs. The linearized modified pET 28a(+) vector was purified with a PCR clean-up kit, and ligated with DNA oligos with T4 DNA ligase at room temperature for 2 hrs. The cells were plated on LB agar plated supplied with 50 μg/mL of kanamycin sulfate. Single *E. coli* colonies were picked, amplified and sequenced to resolve the correct plasmid. Multiple rounds of Pre-RDL cloning were conducted until the gene-of-interest reaches the desired length. In brief, 2 parallel reactions were set up: 4 μg of plasmids encoding ELP genes were either digested by 10 U BseRI and 10 U BglI for 3h, or by 10 U AcuI and 10 U BglI for 3h. Both digestions were separated by low melting point agarose gel, and the correct bands were excised from agarose gel and purified (Omega E.Z.N.A.® Gel Extraction Kit). An equal molar of the parallel reaction products was mixed, and incubated with T4 DNA ligase at room temperature for 1h. The ligation product was transformed into Stbl3 chemically competent cells, amplified and sequenced to resolve ELP constructs with desired lengths. Finally, a short leader sequence (MSKGP) and a short trailer sequence (GWP*) (oligo 21 to oligo 24, table S2) were introduced to the ELP gene through the Pre-RDL method to resolve desired ELP constructs.

Construction of other plasmids:

The gene encoding tau 2N4R, NPM1 and FBL was acquired from the human cDNA library at the Biomedical Research Core Facility at Westlake University. The positively charged C-terminal half of Tau 2N4R between residual 255-441, referred to as tau187, was subcloned into a pET-29b(+) vector using the polymerase incomplete primer extension (PIPE) cloning method (*37*). The NPM1 and FBL were subcloned into the pHTC mammalian expression vector (Promega,

G7711) to generate NPM1-Halo and FBL-Halo plasmids, respectively. The correctness of the insertion for each plasmid was examined by DNA Sanger sequencing.

2. Expression and purification of ELP and tau187 proteins.

ELP purification method was adapted from a previous publication *(22)*. Plasmids were transformed into *E. coli* BL21 DE3* strain (Invitrogen™, C600003). A single colony was picked and inoculated into 5 mL of Terrific Broth (TB) containing corresponding antibiotics and cultured at 37 ˚C with shaking at 220 rpm. When the $OD_{600}$ reached 0.6-0.8, the starting culture was transferred into large culturing flasks containing 1 L TB to allow further growth. When the $OD_{600}$ reached 0.6-0.8, isopropyl ß-D-1-thiogalactopyranoside (Sangon Biotech®, A100487) was introduced to a final concentration of 0.5 mM to induce the expression of ELP. Cultures were allowed to shake at 37˚C for another 16-20 h, harvested and stored at -80˚C freezer. The ELP purification was carried out using the inverse transition cycling (ITC) method (*38*). In brief, the *E. coli* cells were thawed and lysed by sonication in phosphate buffer saline (PBS) in the presence of 1 mM phenylmethyl sulfonyl fluoride (Sangon Biotech®, A610425). The lysate was cleared out by centrifuging at 16,000 g for 60 min at 4 ˚C. To the supernatant, polyethyleneimine (branched, M.W. 1,800, Alfa Aesar) was added to a 0.5 w/v% final concentration to precipitate DNA and RNA, followed by another centrifuging at 16,000 g for 20 min at 4 ˚C to remove DNA and RNA. To the newly obtained supernatant, solid NaCl powder was directly added (12 g NaCl for every 50 mL supernatant, approximately ~4 M NaCl) to precipitate ELP. Once all NaCl was dissolved, the solution was centrifuged at 16,000g for 20 min at 30˚C. The pellet was saved and re-dissolved in ice-cold PBS. A further centrifuge at 16,000g for 20 min at 4˚C was conducted to remove undesired precipitation. Such precipitation-solvation cycle was repeated for another 2 times to improve the purity of ELP. Finally, the ELP was dialyzed against $ddH_2O$ to remove any residual salt. The resulting protein was analyzed by SDS-PAGE to have >95% purity before being concentrated and flash-freeze. The expression and purification of tau187were conducted in accordance with the previously published protocols (*39, 40*).

3. *In-vitro* protein labeling of fluorescent probes using N-Hydroxysuccinimide (NHS) esters and maleimides.

Coumarin-NHS (Sigma-Aldrich®, 36801), fluorescein-NHS (Thermo Scientific™, 46409), fluorescein maleimide (Thermo Scientific™, 62245) and Alexa Fluor™ 647-NHS (Invitrogen™, A37573) were acquired from commercial sources. SBD-NHS, BODIPY-NHS and SBD-maleimide were synthesized in-house and characterized via NMR and mass spectrometry. All fluorophores' stocks were prepared in dry DMSO at 10 mM concentration. For the labeling of NHS ester, ELP (final concentration at ~ 20 mg/mL) and dyes (20% molar ratio of ELP) were mixed in sodium bicarbonate buffer (100 mM $NaHCO_3$, pH 8.4) and allowed to react overnight on ice. The separation of labeled ELP with unreacted dyes was achieved using the PD MidiTrap™ G-25 desalting column (Cytiva®). Tau187 was labeled with fluorescein-maleimide and SBD-maleimide to allow fluorescence intensity and lifetime imaging. For maleimide labeling, tau187 was first exchanged in labeling buffer (20 mM HEPES, pH 7.5, 100 mM NaCl) by dialysis, followed by treatment with 2 mM tris(2-carboxyethyl)phosphine (TCEP) for 1h at room temperature. The labeling reaction was further carried out by adding 100% molar ratio equivalent of dye-maleimide into the reaction mixture, allow to react for 2h on ice. Unreacted dye and TCEP were removed by PD MidiTrap$^{TM}$ G-25 desalting column.

4. <u>Liquid-liquid phase separation (LLPS) assays *in-vitro*.</u>

Phase separation of ELP:

ELPs belong to artificial polypeptides which display lower critical solution temperature (LCST) behavior. The LLPS of ELP could be triggered by the sudden increase of either ionic strength or the environment temperature. Two LLPS conditions were therefore used to initiate ELP LLPS. The first condition is to dilute the concentrated ELPs (originally in $ddH_2O$) into a high-salt (HS) buffer, which results in the final LLPS condition being 70 μM ELP (or 140 μM ELP for the dual-component scenario) in 50 mM HEPES pH 7.0 and 2 M NaCl. Upon mixing thoroughly on ice, the solution was brought to room temperature to trigger the LLPS of ELP in HS condition. Finally, 10 μL of aliquot was transferred onto a glass slide (Sangon Biotech®, F518102) containing a 0.5 mm depth silicon spacer (Grace Bio-Labs, GBL664507). A cover glass (Sangon Biotech®, F518115) was placed on top of the spacer. The entire "sandwich-like" cover glass assembly was flipped over to allow examinations using an inverted confocal microscope. Another LLPS condition was used to examine the phase behavior of ELP in buffers with reduced ionic strength. In this regard, the final LLPS condition was set to be 70 μM ELP (or 140 μM ELP for the dual-component scenario) in 50 mM HEPES pH 7.0 and 500 mM NaCl. Similarly, 10 μL aliquot was transferred into the "sandwich-like" cover glass assembly. The cover glass assembly was heated to 40 ˚C for 10 min on the microscope to initiate LLPS using a built-on heat plate (Instec Inc.).

Phase separation of $V_2I_7E$-40 and $QV_6$-112 in the presence or absence of polyR (in Fig. 3B):

Poly-L-arginine hydrochloride (polyR) consisting of 12 arginines was acquired from a commercial source (Bankpeptide Biological Technology Inc.) and labeled by coumarin-NHS allow fluorescence imaging. 70 μM $V_2I_7E$-40 and 70 μM QV6-112 were mixed in 50 mM HEPES pH 7.0 and 150 mM NaCl in the presence or absence of 50 μM polyR on ice. A low-salt condition was used to avoid the shielding of electrostatic interaction between negative-charged $V_2I_7E$-40 and positive-charged polyR. 10 μL of aliquot was transferred onto a glass slide. A 0.5 mm depth silicon spacer and a cover glass were installed to allow observation under the confocal microscope. The cover glass was heated to 40˚C to initiate LLPS using a built-on heat plate.

Phase separation of tau187 and $V_2I_7E$-40 in the presence or absence of polyU (in Fig. 3C):

Tau187 was first purified and concentrated in the storage buffer containing 20 mM HEPES pH 7.0 and 500 mM NaCl. In the absence of RNA, tau187 remained soluble up to multi-millimolar concentration in this buffer condition. tau187 was mixed with $V_2I_7E$-40 to result in the final LLPS condition to be 70 μM tau187 and 70 μM $V_2I_7E$-40 in 20 mM HEPES pH7.0 and 150 mM NaCl in the absence or presence of 15 μg/mL or 30 μg/mL polyuridylic acid (polyU, 800-1000 kDa, Sigma-Aldrich®, P9528). 10 μL aliquot was transferred to glass slides to allow microscope examination under 40˚C LLPS using a built-on heat plate.

5. Synthetic procedure of SBD and BODIPY-based fluorophores.

All reagents were acquired from commercial sources unless otherwise stated. Reactions were carried out in round bottom flasks (Synthware®) under nitrogen protection and monitored by thin layer chromatography (Synthware®). Products were purified using either flash column chromatography (General-Reagent®) or preparative-scale high-performance liquid chromatography (Waters™ LC Prep 150 System) equipped with reverse phase C18 column (XBridge® Perp C18 5µm OBD™). $^{1}H$ and $^{13}C$ NMR were performed on a Bruker AVANCE NEO 600 MHz spectrometer in $d_6$-dimethylsulfoxide or chloroform-d (Cambridge Isotope Laboratories). High-resolution mass spectra were acquired on Waters SYNAPT-G2-Si ultra-high performance liquid chromatography time-of-flight mass spectrometer. Chemical shifts were reported in reference to the residual solvent peaks of DMSO-$d_6$ or $CDCl_3$. The synthetic route for SBD-NHS ester, SBD-maleimide, SBD-methyl ester and BODIPY-NHS ester could be found in Scheme S1, whereas the synthetic method for S-SBD HaloTag ligand could be found elsewhere *(24)*.

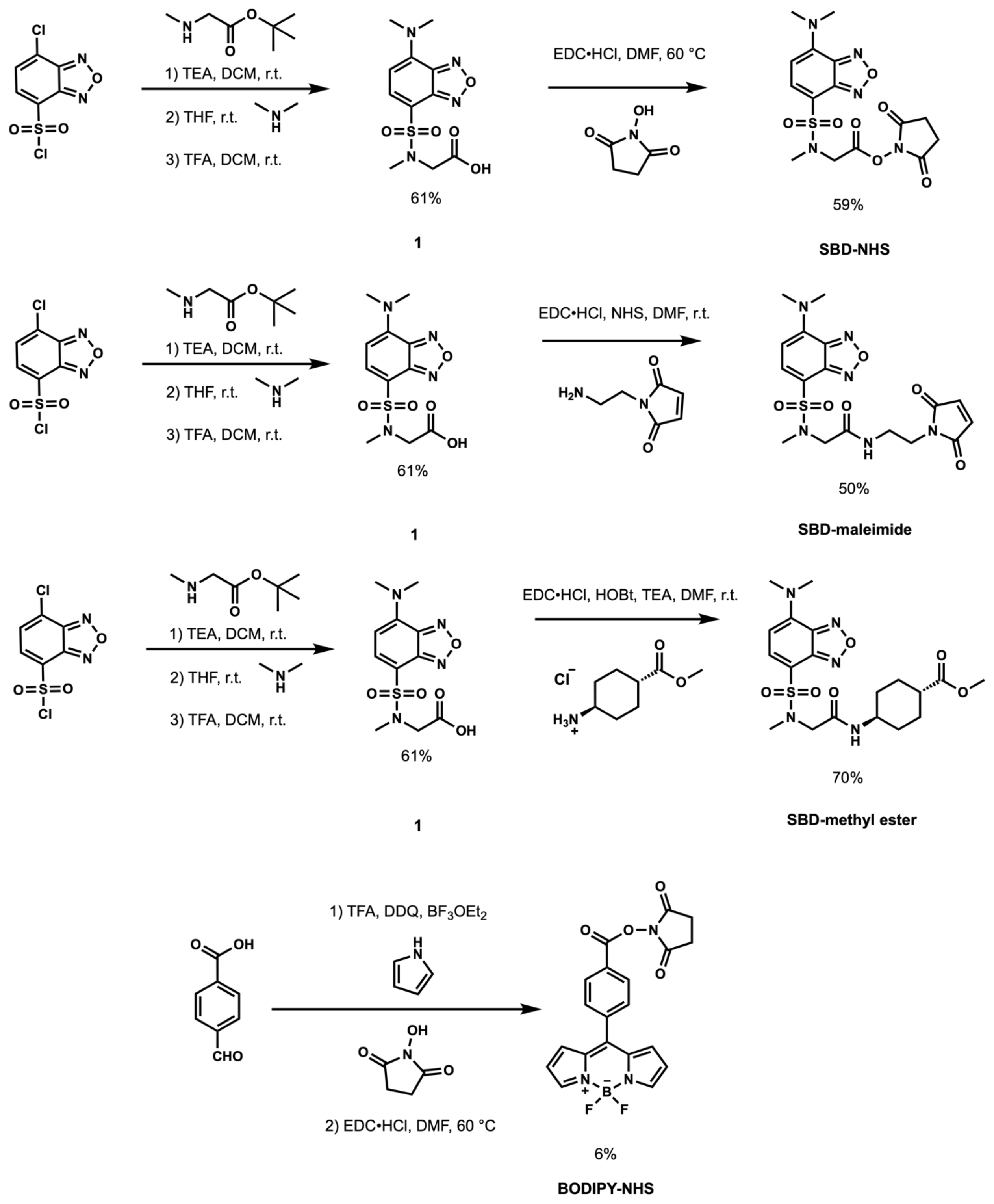


**Scheme S1. Synthetic route for SBD-NHS ester, SBD-maleimide, SBD-methyl ester and BODIPY-NHS ester.**

6. <u>Confocal and fluorescence lifetime imaging (FLIM) imaging procedure for *in-vitro* samples.</u>

The phase-separated samples were first allowed to settle on the glass coverslip for 10 minutes before imaging. Both fluorescence intensity and lifetime images were acquired by a Leica STELLARIS 8 FALCON confocal microscope equipped with a 405 nm laser and a pulsed White Light Laser (WLL) using 63x magnification. For Coumarin, fluorescein and Alexa Fluor™ 647 fluorophores were excited by 405 nm, 504 nm and 653 nm laser lines, respectively. Coumarin and Alexa Fluor™ 647 were excited simultaneously, whereas the excitation of fluorescein was performed using a separate track to avoid crosstalk. Image analysis was achieved using Leica Application Suite X (LAS X) and Fiji software (*41*). FLIM images of SBD and BODIPY labeled *in-vitro* samples were imaged using pulsed laser lines of 448 nm and 488 nm with a repetition rate of 10 and 20 MHz, respectively. Region-of-interest (ROI) was selected using the LAS X FLIM/FCS software to give fluorescence lifetime histograms and the fluorescence lifetime decay curves. Finally, the fluorescence lifetime decay was fitted using a multi-exponential reconvolution model to give intensity-weighted mean fluorescence lifetime ($\tau_{AvInt}$).

7. <u>Mammalian cell culture, confocal and fluorescence lifetime imaging (FLIM) imaging procedure for cell samples.</u>

HEK293T cells were seeded at ~30% confluency in 35 mm glass bottom culture dishes (Cellvis, D35-20-1.5H) treated with poly-L-lysine (Sigma-Aldrich®, P4707) 24 hrs prior to the transfection. Dulbecco's Modified Eagle's Medium (DMEM, Gibco™, 11995-065) was used as the primary culture medium supplemented with 10% Fetal Bovine Serum (FBS, Cellmax, SA211.02, LOT#20210715) and 1x penicillin-streptomycin-glutamine (PSQ, Gibco™, 10378-016). The cells were allowed to grow at 37˚C under 5% $CO_2$ in a Heracell Vios 160i $CO_2$ incubator (Thermo Fisher Scientific).

For FLIM experiments, when cells reach 40% confluency, half of the old DMEM was replaced with fresh DMEM containing 2 μM (2x) S-SBD-Halo. Hence a final 1 μM S-SBD-Halo fluorophore was used to allow FLIM imaging. Additionally, 1 μg mammalian expression plasmid containing gene-of-interest with HaloTag was transiently transfected into HEK293T cells using 3 μL X-tremeGENE 9 DNA transfection reagent (Roche) diluted in 100 μL 1x OPTI-MEM® I reduced serum medium (Gibco™, 31985-062). The cells were put back into the incubator at 37˚C under 5% $CO_2$ to allow protein expression and labeling for 24 hrs. After 24 hrs, the cells were washed with fresh, colorless DMEM/F12 (1:1) medium (Gibco™, 21041-025) containing 10% FBS to remove unbounded HaloTag substrate and phenol red. Live cell FLIM experiments were achieved inside a controlled environmental chamber (37˚C, 5% $CO_2$, Okolab) built on a Leica STELLARIS 8 FALCON confocal microscope equipped with a pulsed WLL setting at 448 nm with a 10 MHz repetition rate. For the demonstration of the nucleolar cap, additional cell samples were treated with actinomycin D (Act D, Gibco™, 11805-017) with a final concentration of 0.2 μM 2 hrs prior to the FLIM imaging. FLIM data analysis was achieved using the LAS X FLIM/FCS software.

For dual-color confocal imaging experiments, half of the DMEM was replaced with fresh DMEM when cells reach 40% confluency in 35 mm glass bottom dishes. A final 0.5 μM TMR HaloTag® ligand (Promega, G8251) was added to the medium to specific labeling with FBL-Halo to allow confocal imaging. 1 μg plasmid encoding EGFP-NPM1 and 1 μg plasmid encoding FBL-Halo were co-transfected into HEK293T cells using 6 μL X-tremeGENE 9 DNA transfection reagent diluted in 200 μL 1x OPTI-MEM® I reduced serum medium. After 24 hrs of expression, Hoechst 33342 (Invitrogen™, H1399) was added to the cell culture with a final concentration of

5 μg/mL. The cells were further incubated for 10 minutes at a 37˚C incubator before a quick PBS wash to remove excess fluorophores. The cells were fixed with 4% formaldehyde solution (Thermo Scientific$^{TM}$, 28908) for 10 minutes and washed with PBS. The coverslip was mounted with sterile 50% glycerol. Hoechst 33342, EGFP and TMR were excited by 405 nm, 488 nm and 550 nm laser lines, respectively.

8. Simulation details for ELP dielectric constant and surface tension.

Simulation details:

To investigate the relationship between surface tension and dielectric constant in biological condensates, we performed multiscale simulations on ELP condensates. Specifically, we examined ten ELP repeat versions for five of the sequences studied experimentally, namely $V_{10}$, $V_5A_2G_3$, $QV_6QV_2$, $V_2I_7E$, and $KV_6KV_2$. For charged amino acids K and E, both charged ($V_2I_7E^-$ and $K^+V_6K^+V_2$) and uncharged ($V_2I_7E^0$ and $K^0V_6K^0V_2$) versions were simulated. All simulations were performed in 1 M NaCl. Our simulation methodology began with homopolymer and MOFF simulations (*42-44*) to generate initial structures. Then, we utilized simulations in MARTINI3 *(35)* to report on condensate surface tension. Finally, we probed the dielectric constant of condensates with all-atom simulations in CHARMM36m with shifted water interactions to correct for disordered proteins *(36)*.

To ensure the protein density was the same for each condensate, we began with homopolymer simulations. Our homopolymer force field was based on the MOFF force field, with random coil secondary structure and non-bonded interactions identical to MOFF's valine-valine interaction. In this force field, we placed 40 homopolymers of 50 residues in length inside a 75 nm by 75 nm by 75 nm simulation box. After the steepest decent energy minimization, we performed an NPT simulation for 0.1 μs at 150 K and 1 bar, resulting in a dense polymer phase.

To optimize specific protein interactions within this phase, we performed MOFF simulations (*42-44*). As ELPs are known to have minimal secondary structures in both the dense and dilute state (*45, 46*), we assumed the secondary structure to be similar to that of a random coil and set the equilibrium angle between residues to 127˚ in all cases. In this force field, we conducted energy minimization, followed by a 0.1 μs NVT simulation, with a timestep of 10 fs. During the simulation, we linearly increased the temperature from 150 K to 300 K. Nonbonded parameters for neutral K and E were identical to the charged versions, but with the charge changed to 0.

The final snapshots from MOFF were used as a starting point for explicit solvent coarse-grained simulations in MARTINI. MOFF is a one-bead per amino acid representation, so we converted this simplified snapshot to an all-atom representation using tleap, which is part of AMBER tools 2020 (*47*). We then converted the all-atom configuration to a MARTINI3 configuration using martinize2 (*43*). We enforced neutral termini of the peptide chains and used random coil secondary structure throughout the entire peptide. Next, we expanded the Z-dimension of the simulation box to 40 nm and centered the protein in the simulation box. Thus, all simulations started with box dimensions of 6.3782 nm by 6.3782 nm by 40.0000 nm, which allowed for slab simulations to measure the coexistence between the dense phase and the dilute phase (*48*). After solvating with MARTINI water and 1 M NaCl, we performed the steepest decent energy minimization. Then, we performed NVT equilibration at 313.15 K for 5 ns with a timestep of 10 fs. Next, we performed NPT equilibration at 1 bar and 313.15 K for 10 ns with a timestep of 20 fs. We then used semi-isotropic pressure coupling, sometimes called the $NP_NAT$ ensemble, to perform production simulations for 100 μs and used a timestep of 20 fs. The initial 25 μs was discarded for equilibration, leaving 75 μs for analysis. The temperature of these simulations was held at 313.15 K, while the pressure normal to the surface was fixed at 1 bar. This setup allowed for the calculation of surface tension, as discussed below (*49*).

Dielectric constant calculations were based on the fluctuations of partial charges, and thus required all-atom simulations. These simulations were performed in the CHARMM36m force field. Proteins were capped using an acetylated N-terminus and an amidated C-terminus. TIP3P water was used, and the water-hydrogen Lennard-Jones well depth was scaled to -0.10 kcal/mol to better describe IDPs, as has been done previously *(36)*. To ensure the reproducibility of our results, we converted 3 different MARTINI configurations for each protein into starting configurations for all-atom simulations. To begin these all-atom simulations, MARTINI simulations at 50 μs, 75 μs, and 100 μs were converted into the CHARMM36m representation using the backward script (*50*). From the output of the backward calculation, we performed energy minimization, followed by an NVT simulation at 313.15 K for 100 ps with a timestep of 1 fs and with position restraints on both the protein backbone and side chain. Next, we performed NPT equilibration at 313.15 K and 1 bar for 5 ns with a timestep of 2 fs. We then conducted $NP_NAT$ at 313.15 K and 1 bar in the Z-dimension for 5 ns with a timestep of 2 fs. Finally, we performed $NP_NAT$ production runs for 250 ns with a timestep of 2 fs at 313.15 K and 1 bar in the Z-dimension. The first 50 ns was discarded for equilibration, leaving 200ns for analysis.

Finally, not every coarse-grained configuration can be stably converted into higher resolution. When the final snapshot from MOFF simulations failed to result in stable MARTINI representations or the final snapshot from MARTINI simulations failed to result in stable all-atom representations, we iteratively moved back one snapshot and repeated the conversion procedure until a stable configuration was found.

Simulation analysis：

All simulations were analyzed to determine the number of protein clusters. Our code first computed a center of mass contact matrix between peptide molecules with a distance cutoff of 4 nm. Then, using a depth-first search algorithm (*51*), we identified the size of each cluster. Our code was implemented in MDanalysis (*52*).

Our simulation layout placed a dense slab in the x-y plane, and the solvent created a coexisting phase in the z-dimension. In this configuration, we controlled the pressure normal to the protein-solvent interface by performing simulations in the $NP_NAT$ ensemble. These assumptions allowed us to calculate the surface tension (τ) as

$$\tau = \frac{L_Z}{2N} * (P_{ZZ} - \frac{1}{2}(P_{XX} + P_{YY}))$$

where $L_Z$ is the length of the Z-dimension, $P_{ZZ}$ is the Z-component of the component of the pressure tensor, $P_{XX}$ is the X-component of the component of the pressure tensor, and $P_{YY}$ is the Y-component of the component of the pressure tensor. We extracted the number of droplets by using our clustering algorithm described above, and $N$ is the number of clusters containing 10 or more peptides. Error bars represent the standard deviation over five independent time windows.

To compute the dielectric constant of each condensate, we first centered our simulation on the center of mass for proteins that remained in the largest cluster for over 99% of the simulation time. Next, we computed the dielectric in 20 slabs across the simulation box as

$$\varepsilon_0 = 1 + \frac{4\pi}{3\bar{V}k_BT}(< M^2 > - < M >^2)$$

where $\varepsilon_0$ is the dielectric constant, $\bar{V}$ is the average volume, $k_B$ is Boltzmann's constant, $T$ is temperature, and $M$ is the dipole moment of the slab. For each slab, we extracted the dielectric constant at the final timepoint, and constructed a plot of the dielectric constant as a function of Z-axis of the simulation box. We then reported the final dielectric constant by sorting the dielectric profile from the lowest to the highest dielectric, and reporting the histogram bin with the lowest dielectric constant, which corresponds to the region of the simulation with the highest protein density and thus is likely the best approximation for the interior of the condensate.

### Characterization

**Compound 1.** N-((7-(dimethylamino)benzo[c][1,2,5]oxadiazol-4-yl)sulfonyl)-N-methylglycine. $^1$H NMR (500 MHz, $CDCl_3$) δ 7.91 (d, *J* = 8.3 Hz, 1H), 6.03 (d, *J* = 8.3 Hz, 1H), 4.26 (s, 2H), 3.52 (s, 6H), 2.94 (s, 3H). $^{13}$C NMR (151 MHz, DMSO) δ 170.14, 146.72, 144.89, 142.93, 138.32, 107.64, 101.57, 50.58, 42.29, 35.39. $[M+Na]^+$ Calcd, 337.0583, Obsd, 337.0582.

**SBD-NHS ester.** 2,5-dioxopyrrolidin-1-yl N-((7-(dimethylamino)benzo[c][1,2,5]oxadiazol-4-yl)sulfonyl)-N-methylglycinate.
$^1$H NMR (500 MHz, $CDCl_3$) δ 7.89 (d, *J* = 8.3 Hz, 1H), 6.02 (d, *J* = 8.3 Hz, 1H), 4.59 (s, 2H), 3.50 (s, 6H), 3.01 (s, 3H), 2.80 (s, 4H). $^{13}$C NMR (126 MHz, $CDCl_3$) δ 168.56, 164.83, 146.78, 145.18, 143.74, 138.61, 108.94, 101.31, 49.49, 42.78, 35.61, 25.66. $[M+H]^+$ Calcd, 412.0927, Obsd, 412.0929.

**SBD-maleimide.** 2-((7-(dimethylamino)benzo[c][1,2,5]oxadiazole)-4-sulfonamido)-N-(2-(2,5-dioxo-2,5-dihydro-1H-pyrrol-1-yl)ethyl)acetamide.
$^1$H NMR (600 MHz, $CDCl_3$) δ 7.95 (d, *J* = 8.4 Hz, 1H), 6.69 (s, 2H), 6.06 (d, *J* = 8.4 Hz, 1H), 3.86 (s, 2H), 3.79-3.70 (m, 2H), 3.59-3.50 (m, 8H), 2.82 (s, 3H). $^{13}$C NMR (151 MHz, $CDCl_3$) δ 170.76, 168.91, 146.60, 145.11, 144.01, 139.98, 134.19, 106.38, 101.20, 53.30, 42.79, 38.47, 37.53, 36.66. $[M+Na]^+$ Calcd, 459.1063, Obsd, 459.1071.

**SBD-methyl ester.** Methyl-4-(2-((7-(dimethylamino)-benzo[c][1,2,5]oxadiazole)-4-sulfonamido)acetamido)cyclohexane-1-carboxylate.
$^1$H NMR (500 MHz, $CDCl_3$) δ 7.95 (d, *J* = 8.3 Hz, 1H), 6.95 (d, *J* = 8.0 Hz, 1H), 6.06 (d, *J* = 8.4 Hz, 1H), 3.88 (s, 2H), 3.82-3.73 (m, 1H), 3.68 (s, 3H), 3.54 (s, 6H), 2.81 (s, 3H), 2.35-2.25 (m, 1H), 2.09-2.01 (m, 4H), 1.61-1.53 (m, 2H), 1.32-1.24 (m, 2H). $^{13}$C NMR (151 MHz, $CDCl_3$) δ 175.89, 167.44, 146.66, 145.21, 144.16, 140.13, 106.18, 101.30, 53.44, 51.77, 47.91, 42.92, 42.40, 36.56, 31.99, 27.87. $[M+H]^+$ Calcd, 454.1760, Obsd, 454.1757.

**S-SBD-Halo.** N-(2-(2-((6-chlorohexyl)oxy)ethoxy)ethyl)-4-(2-((7-(dimethylamino)-benzo[c][1,2,5]thiadiazole)-4-sulfonamido)acetamido)cyclohexane-1-carboxamide.
$^1$H NMR (600 MHz, DMSO) δ 7.97 (s, 1H), 7.76-7.68 (m, 2H), 6.55 (s, 1H), 3.82 (s, 2H), 3.64-3.59 (m, 2H), 3.48-3.44 (m, 9H), 3.32-3.30 (m, 4H), 3.19-3.13 (m, 2H), 2.77 (s, 3H), 2.07-1.98 (m, 1H), 1.76-1.64 (m, 7H), 1.52-1.44 (m, 2H), 1.41-1.26 (m, 7H), 1.18 – 1.08 (m, 2H). $^{13}$C NMR (151 MHz, $CDCl_3$) δ 175.16, 167.91, 152.36, 147.85, 147.36, 137.03, 112.79, 103.46, 71.39, 70.37, 70.12, 69.92, 54.36, 47.93, 45.18, 44.61, 42.99, 39.13, 36.85, 32.63, 32.36, 29.57, 28.49, 26.79, 25.52. $[M+H]^+$ Calcd, 661.2609, Obsd, 661.2607.

**BODIPY-NHS ester.** 2,5-dioxopyrrolidin-1-yl 4-(5,5-difluoro-5H-4λ4,5λ4-dipyrrolo[1,2-c:2',1'-f][1,3,2]diazaborinin-10-yl)benzoate.
$^1$H NMR (600 MHz, $CDCl_3$) δ 8.31 (d, *J* = 8.4 Hz, 2H), 7.99 (s, 2H), 7.72 (d, *J* = 8.4 Hz, 2H), 6.86 (d, *J* = 3.6 Hz, 2H), 6.65-6.55 (m, 2H), 2.96 (s, 4H). $^{13}$C NMR (151 MHz, $CDCl_3$) δ 169.18, 161.30, 146.66, 145.40, 144.88, 140.09, 134.76, 131.45, 130.85, 130.74, 127.22, 119.39, 25.85. $[M+H]^+$ Calcd, 410.1127, Obsd, 410.1127.

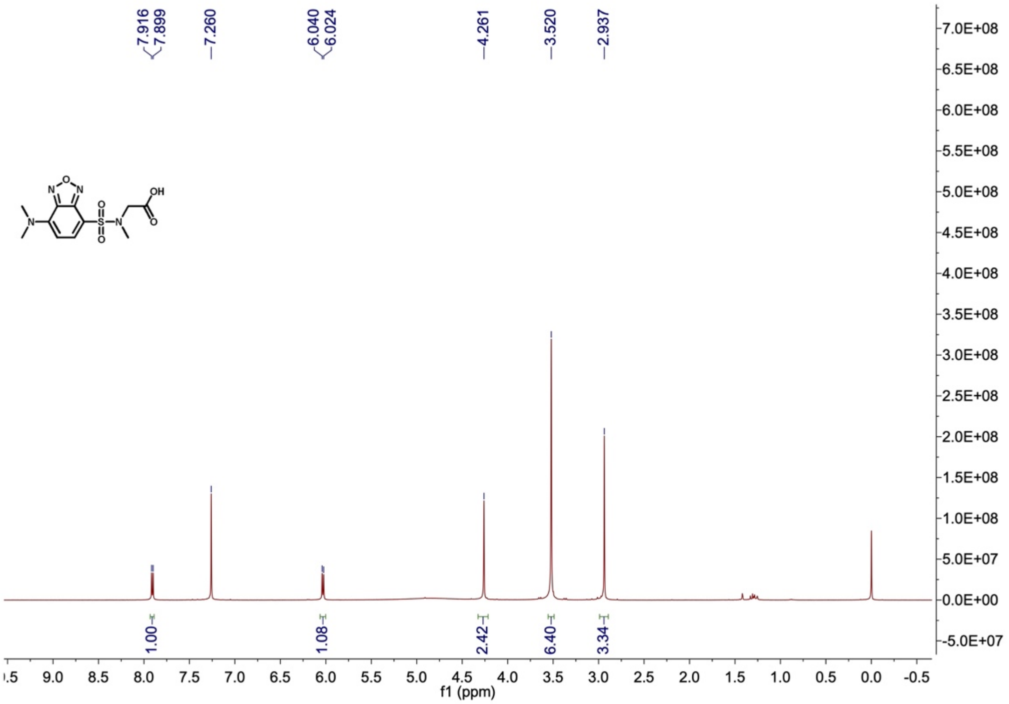


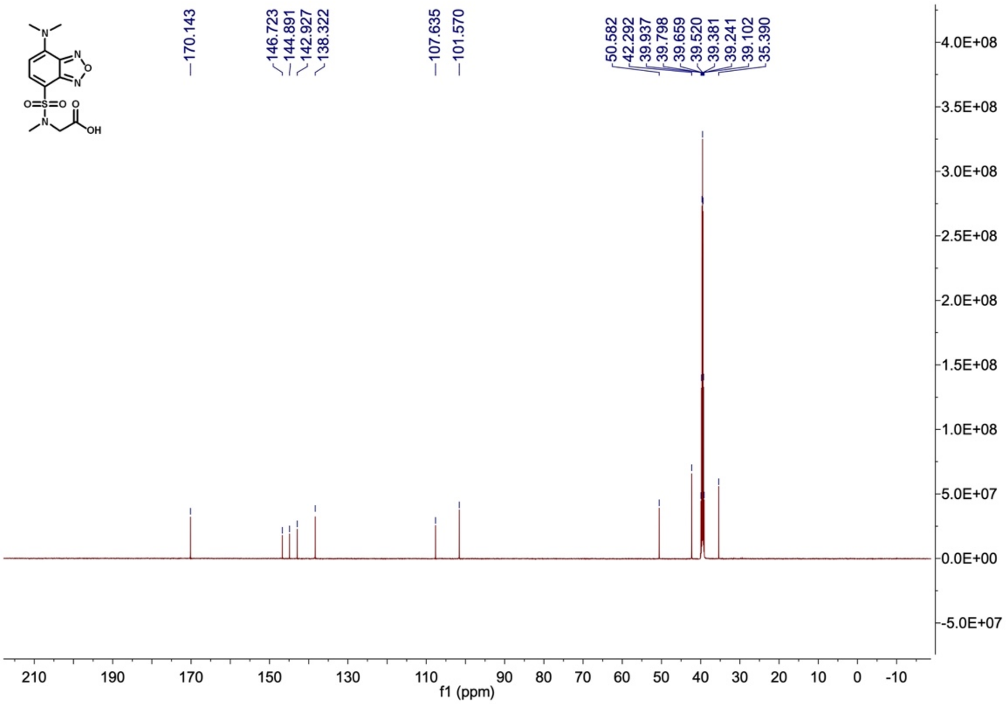


$^1H$ and $^{13}C$ NMR spectra for compound **1**.

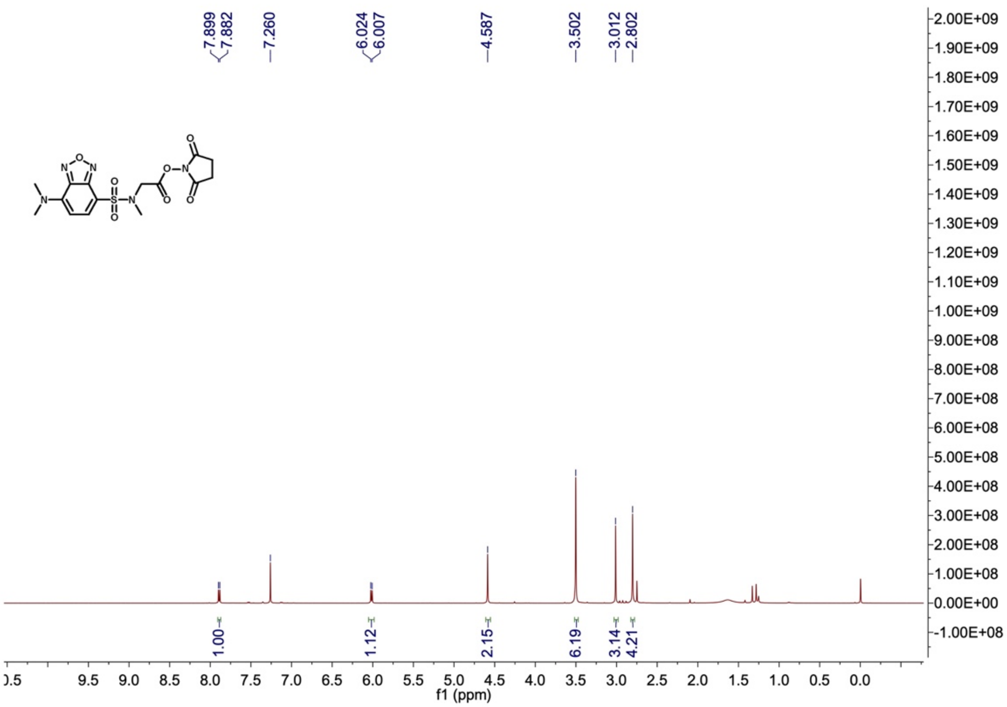


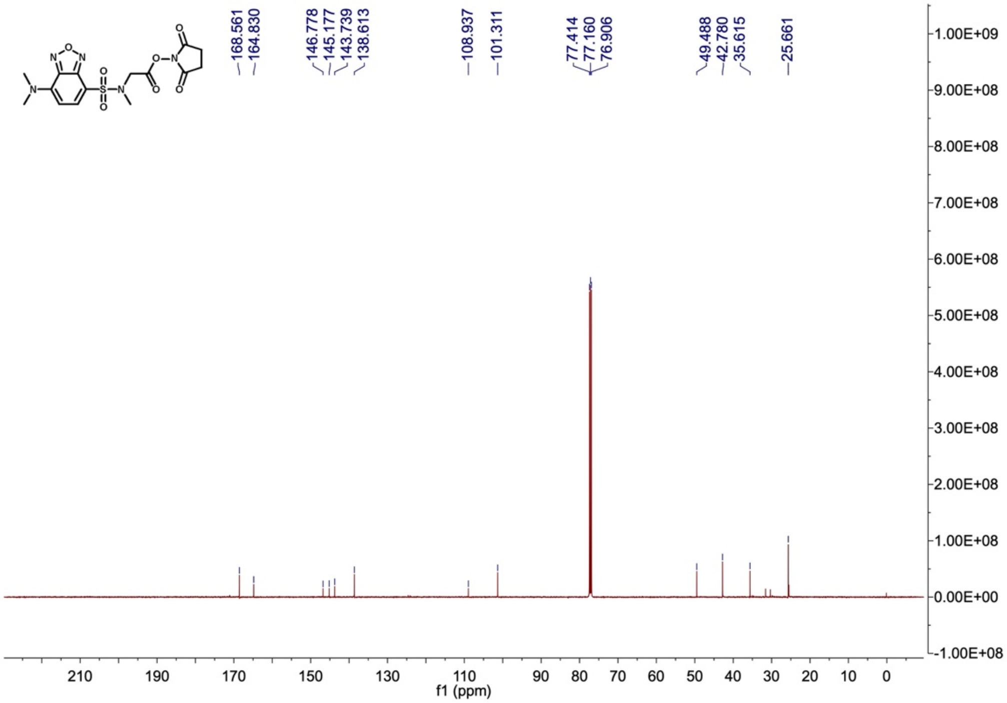


$^{1}$H and $^{13}$C NMR spectra for **SBD-NHS.**

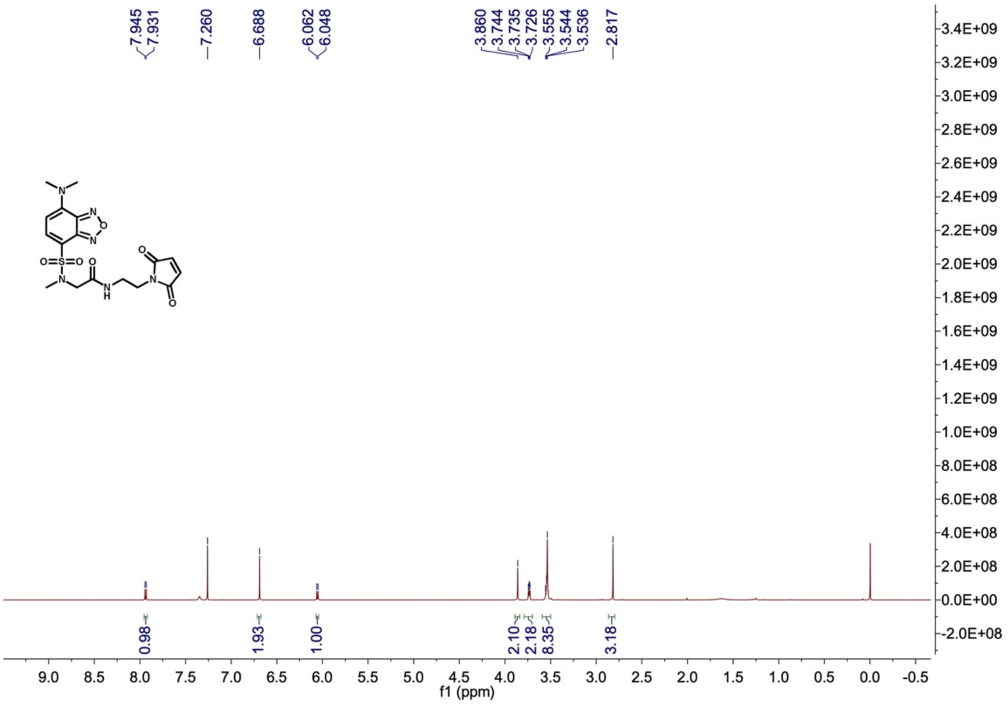


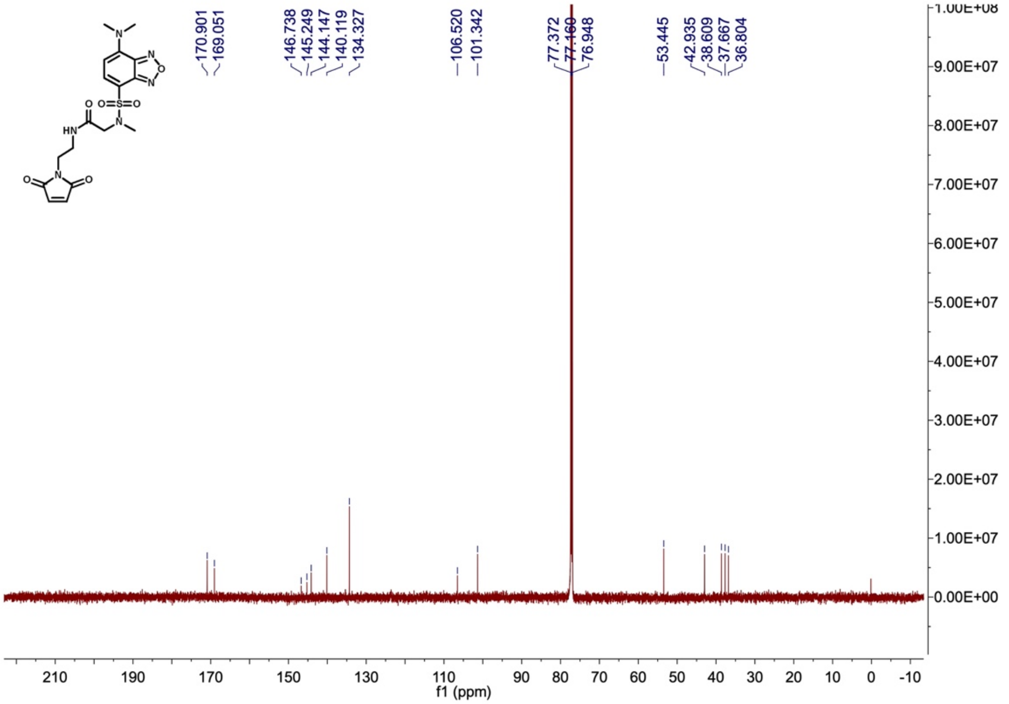


$^{1}H$ and $^{13}C$ NMR spectra for **SBD-maleimide**.

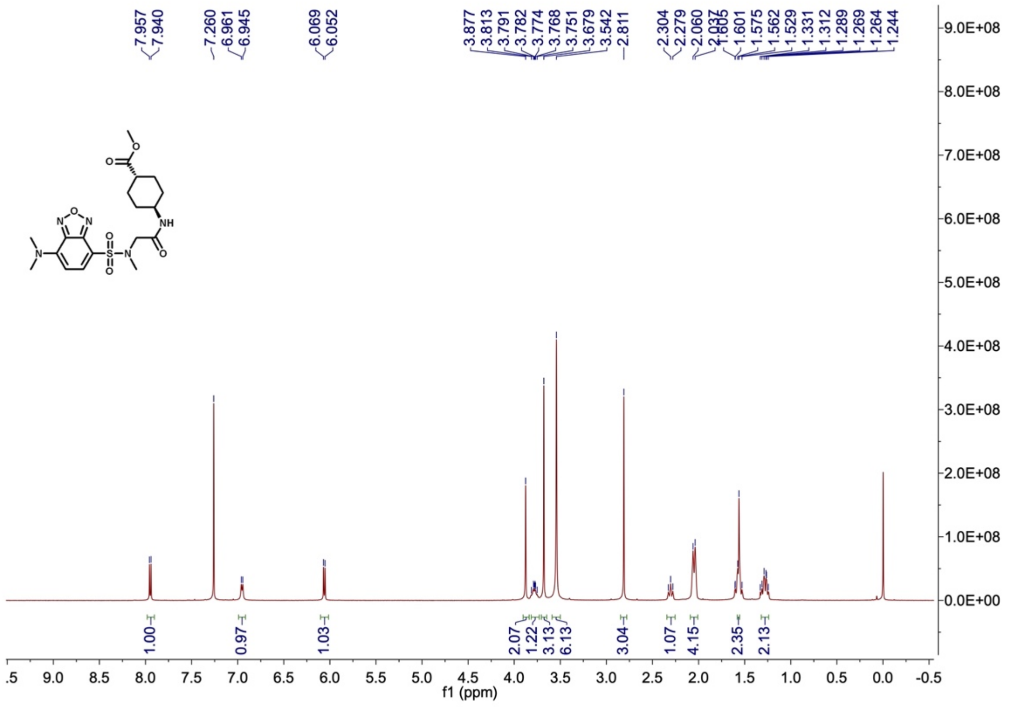

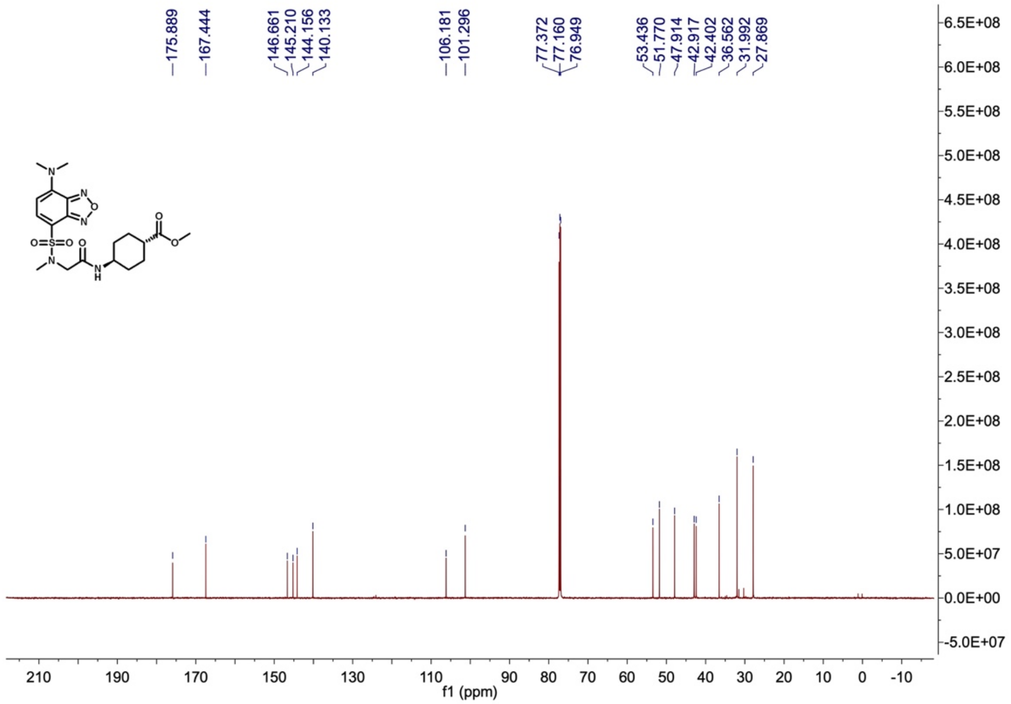

$^{1}H$ and $^{13}C$ NMR spectra for **SBD-methyl ester**.

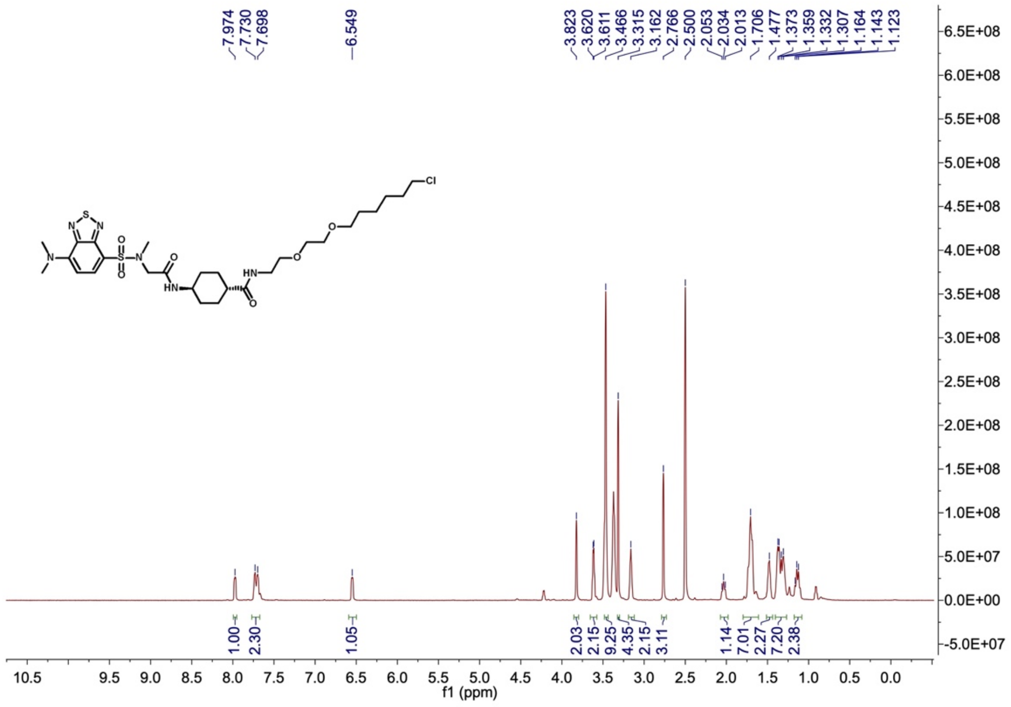

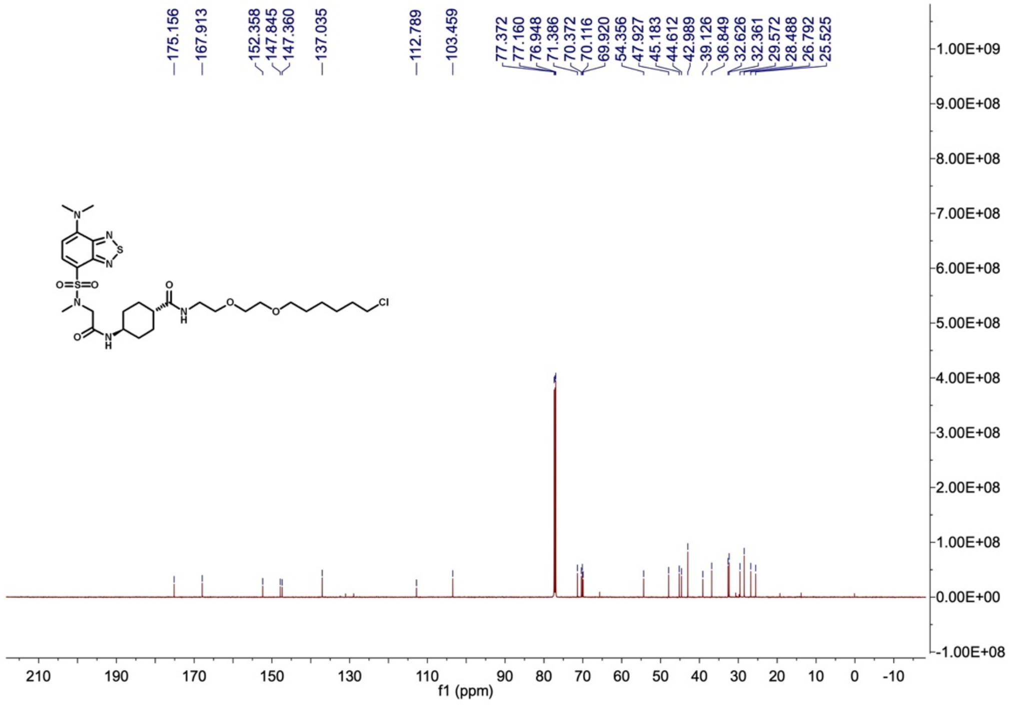

[1]H and [13]C NMR spectra for **S-SBD-Halo**.

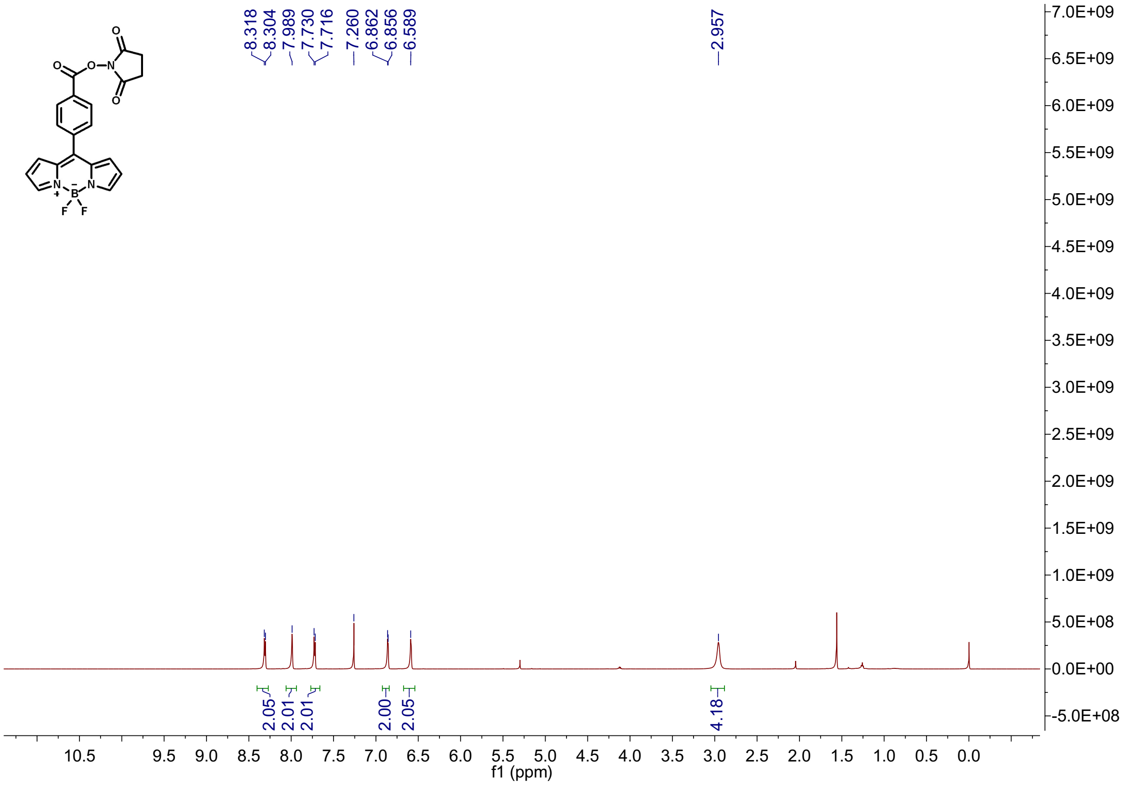


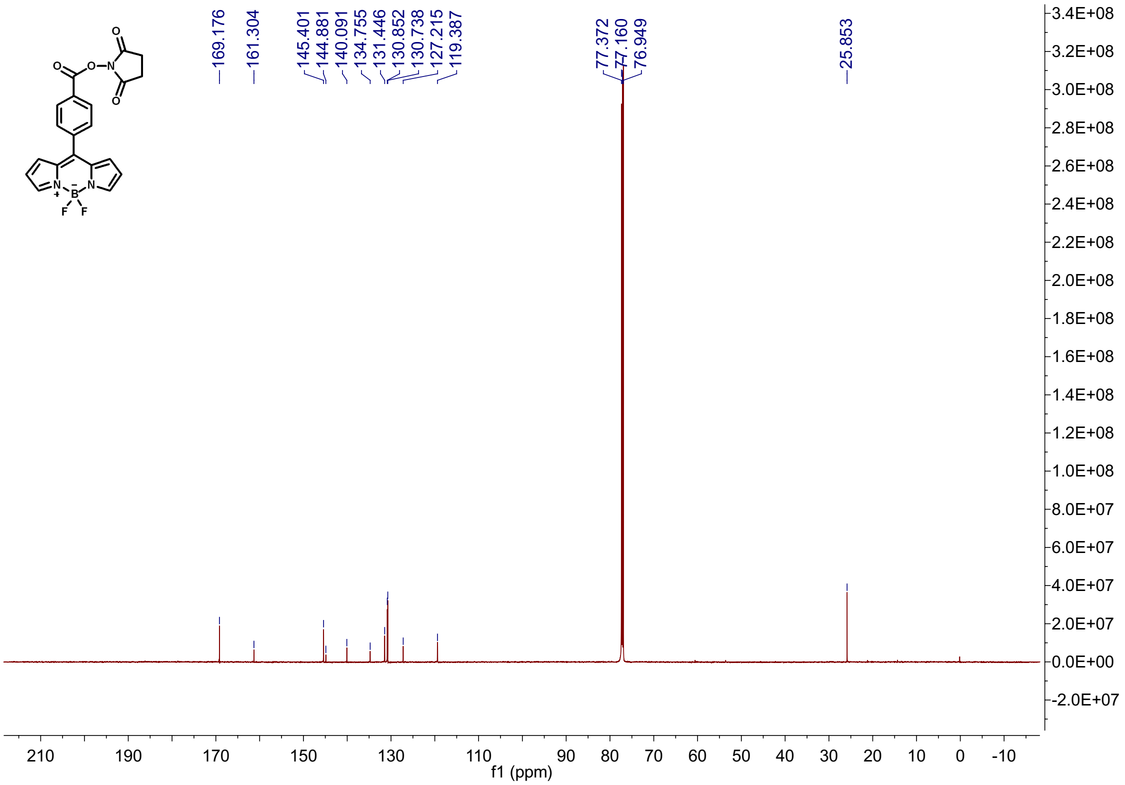


$^{1}H$ and $^{13}C$ NMR spectra for **BODIPY-NHS**.

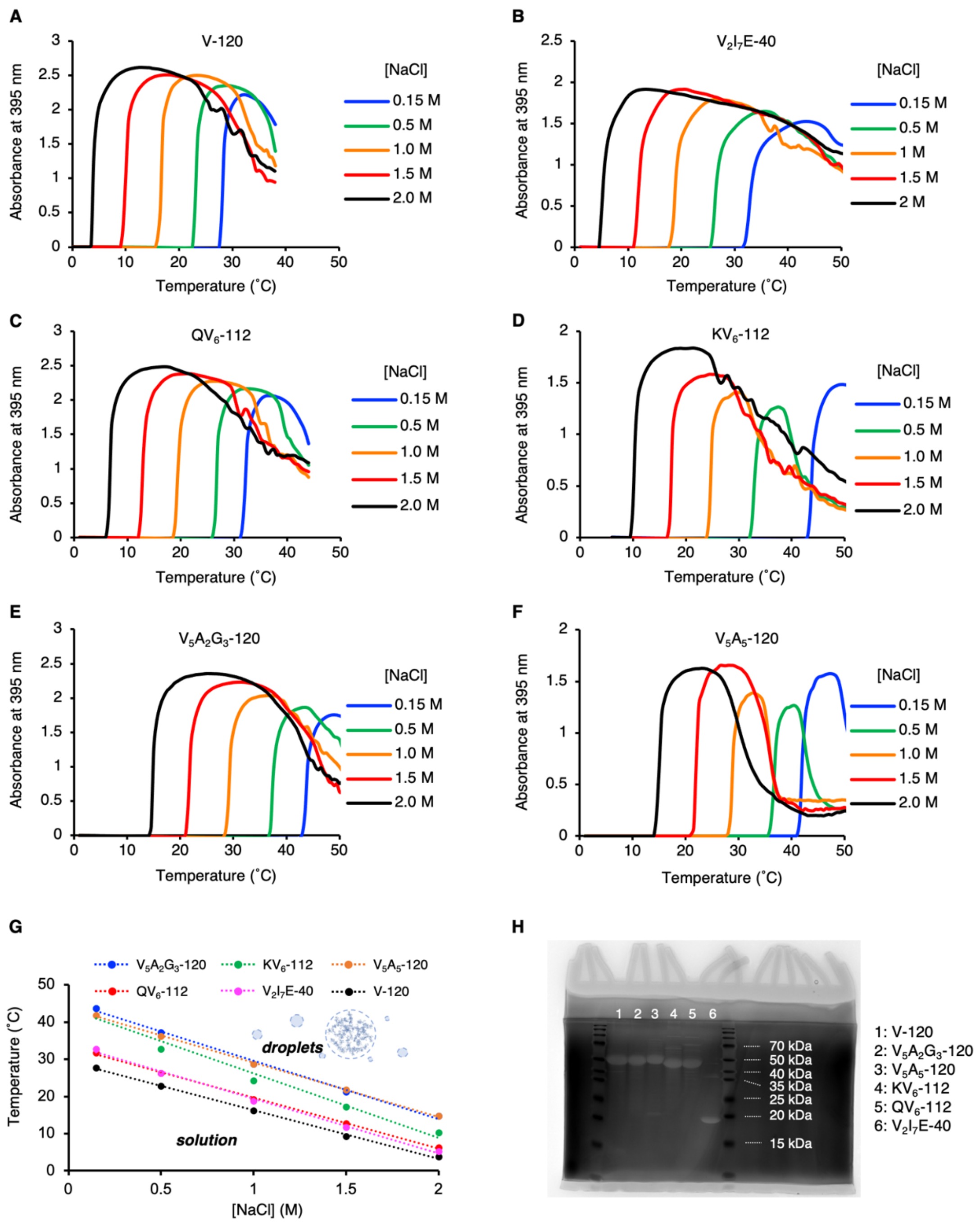


**Fig. S1. Analyzation of the phase transition temperature and purity of ELP. (A-F)** Phase transition processes of ELP were captured by the rapid increase of absorbance signal using a UV-Vis-NIR spectrometer. ELPs were mixed in buffers (50 mM HEPES, pH 7.0) with different molarity of NaCl (0.15 M, 0.5 M, 1.0 M, 1.5 M, 2.0 M) at a final peptide concentration of 70 µM on ice. An assay was designed to gradually increase the sample temperature from 1 ˚C to 50 ˚C at 0.5 ˚C/min rate, while the absorbance at 395 nm was recorded for every 0.5˚C of temperature increment. **(G)** Summary of the phase transition temperature of ELPs at different NaCl concentrations. **(H)** SDS-PAGE analysis of the purity of the ELP. The polyacrylamide gel was stained in 0.5 M $CuCl_2$ solution to develop protein bands.

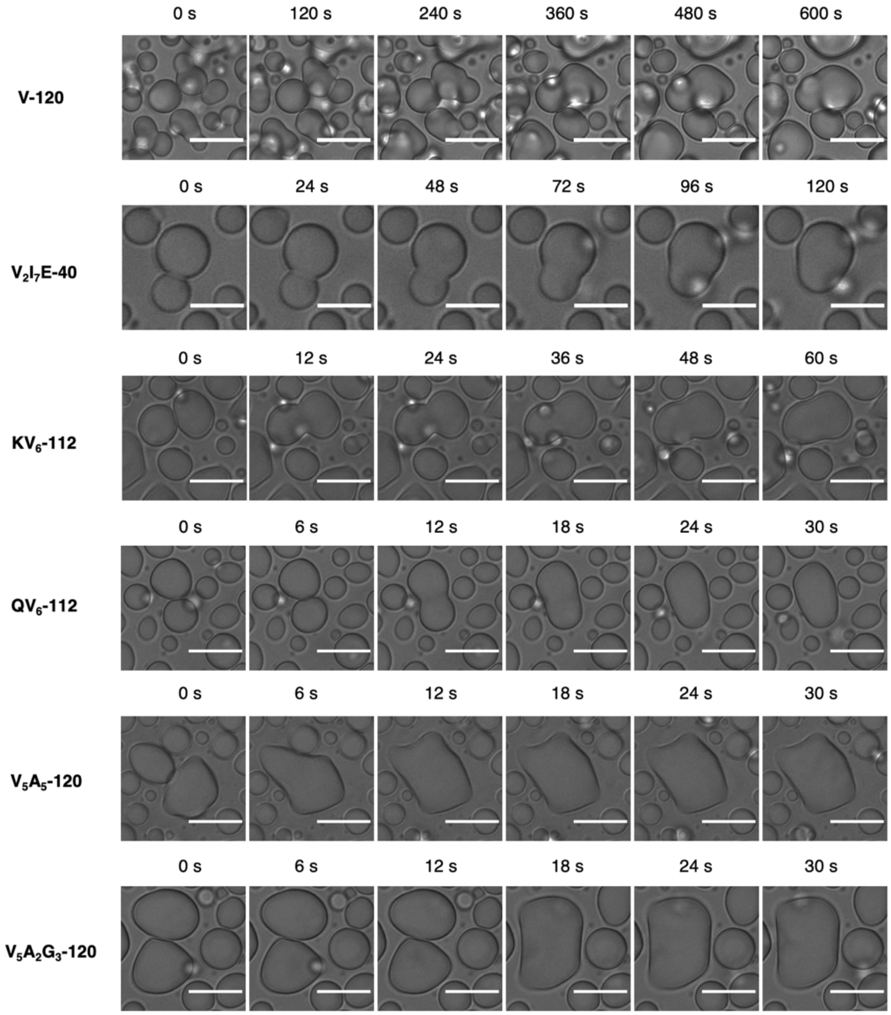


**Fig. S2. ELP condensates are liquid-like droplets displaying extensive wetting and merging behavior.** Time-lapse imaging was conducted with 70 µM ELP in phase separation buffer (50 mM HEPES, pH 7.0, 2 M NaCl) at room temperature. Scale bar for $V_2I_7E$-40, 5 µm. Scale bars for other ELPs, 10 µm.

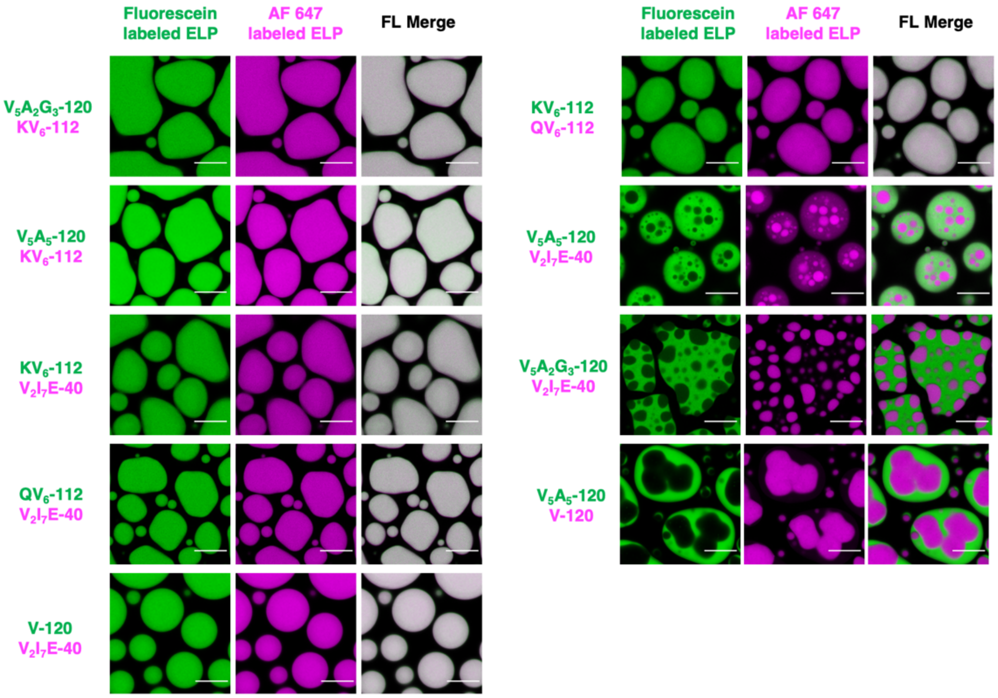


**Fig. S3. Confocal images of dual-component ELP condensates demonstrated diverse phase behavior.** Co-phase separation of 70 µM fluorescein-labeled ELP (green) and 70 µM AF 647-labeled ELP (magenta) in phase separation buffer (50 mM HEPES, pH 7.0, 2 M NaCl) at room temperature resulted in both miscible ELP condensates (left column), partially miscible and core-shell type dual-layer ELP droplets (right column). Scale bars, 10 µm.

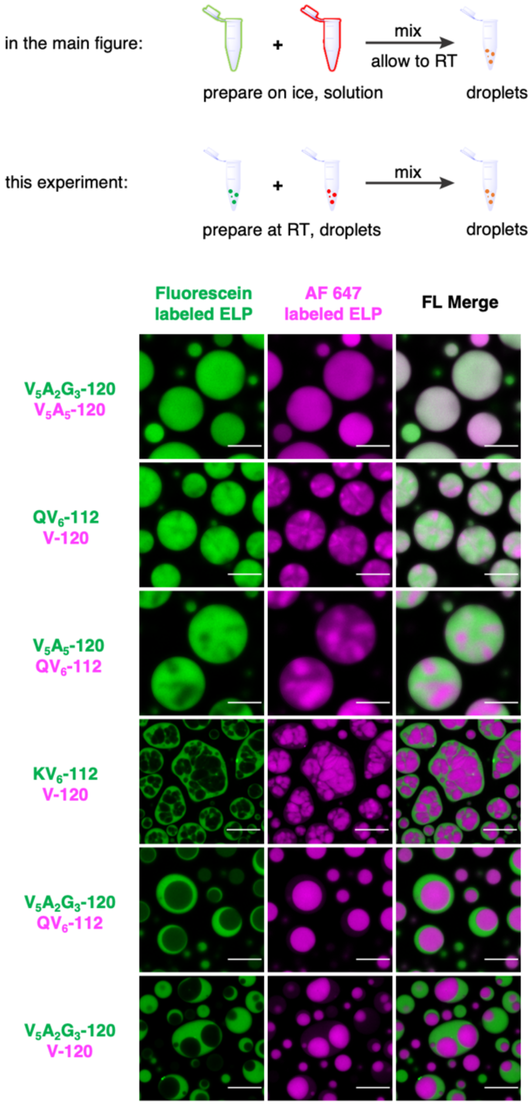


**Figure S4. Dual component ELP droplets prepared by mixing two already phase-separated ELP samples resulted in identical core-shell structures compared to the co-phase separation of binary ELP solution mixtures.** Two different ELPs (labeled with fluorescein or AF 647) were allowed to phase separate individually in phase separation buffer (50 mM HEPES, pH 7.0, 2 M NaCl) at a final concentration of 140 µM at room temperature. The two phase-separated samples were then mixed with an equal amount and allowed to settle for 15 min on glass coverslips for confocal imaging. Scale bars, 10 µm.

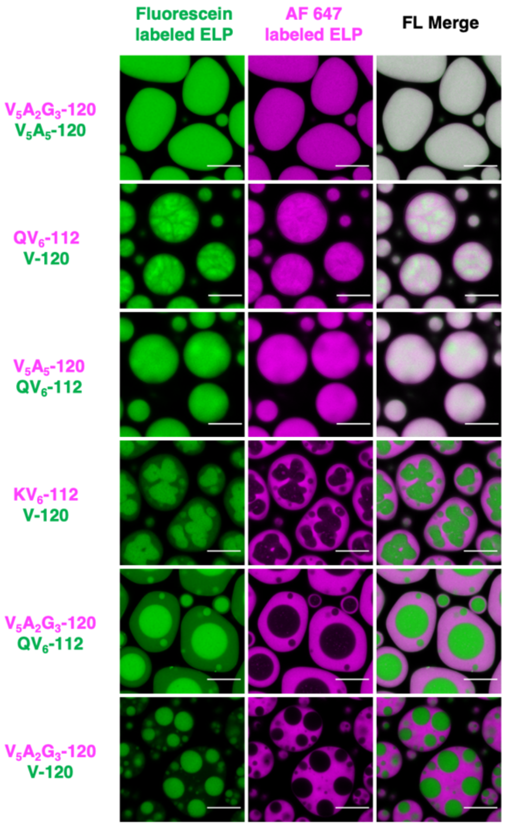


**Figure S5. Swapping fluorophores on ELP preserved the organization of dual-component ELP condensates.** Repeating dual color imaging from **Fig 1C** with swapped labeling fluorophores yield the dual component ELP condensates with identical organizations but inverted colors. Scale bars, 10 µm.

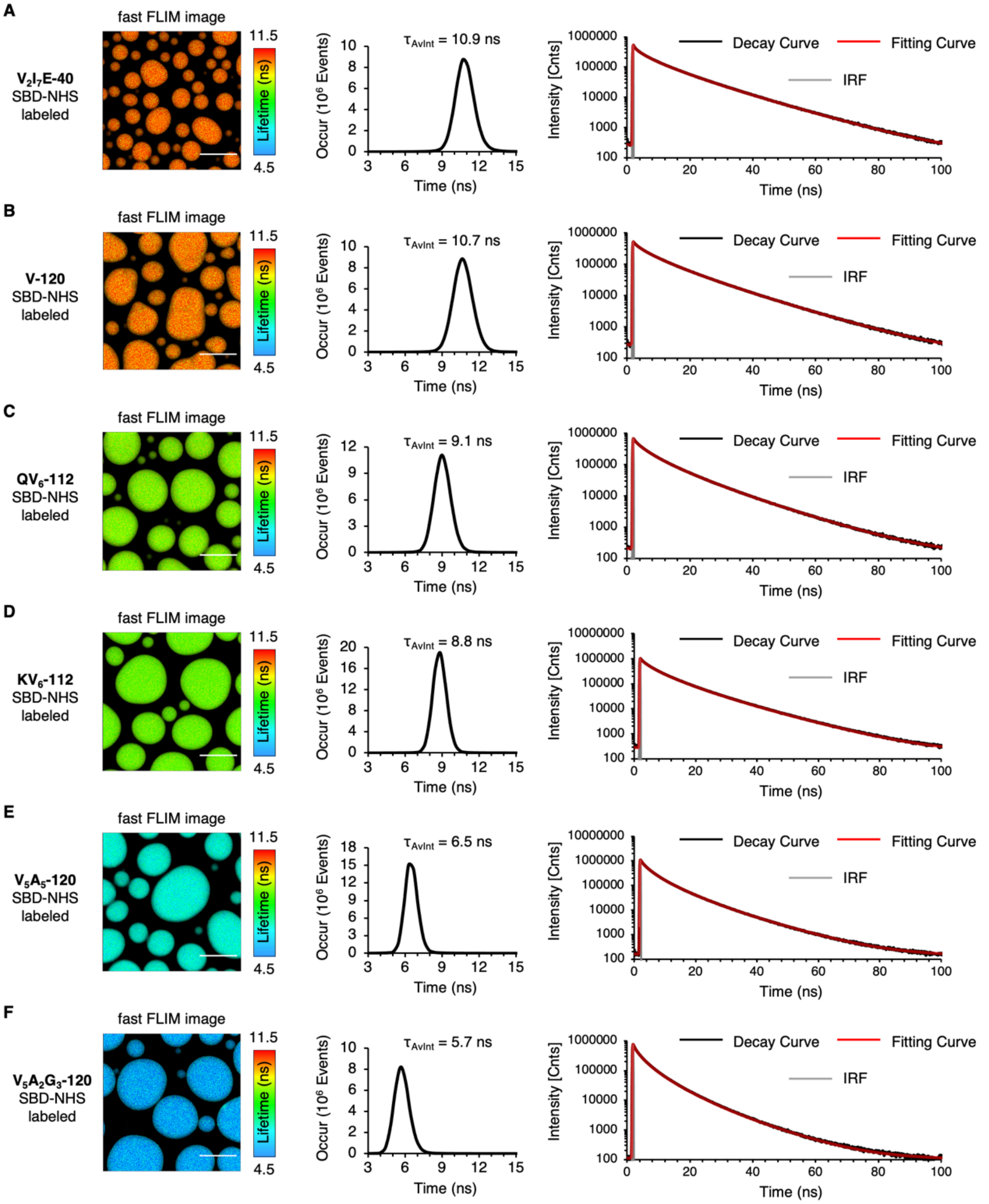


**Figure S6. Photophysical data of SBD-NHS labeled ELP FLIM measurements in 2 M NaCl solution. (A-F)** Fast FLIM images, fluorescence lifetime histograms and fluorescence lifetime decay curves of SBD-labeled ELP condensates in 2 M NaCl solution. 70 µM ELP labeled with SBD-NHS were mixed in phase separation buffer (50 mM HEPES, pH 7.0, 2 M NaCl) to allow FLIM imaging. Scale bars, 10 µm.

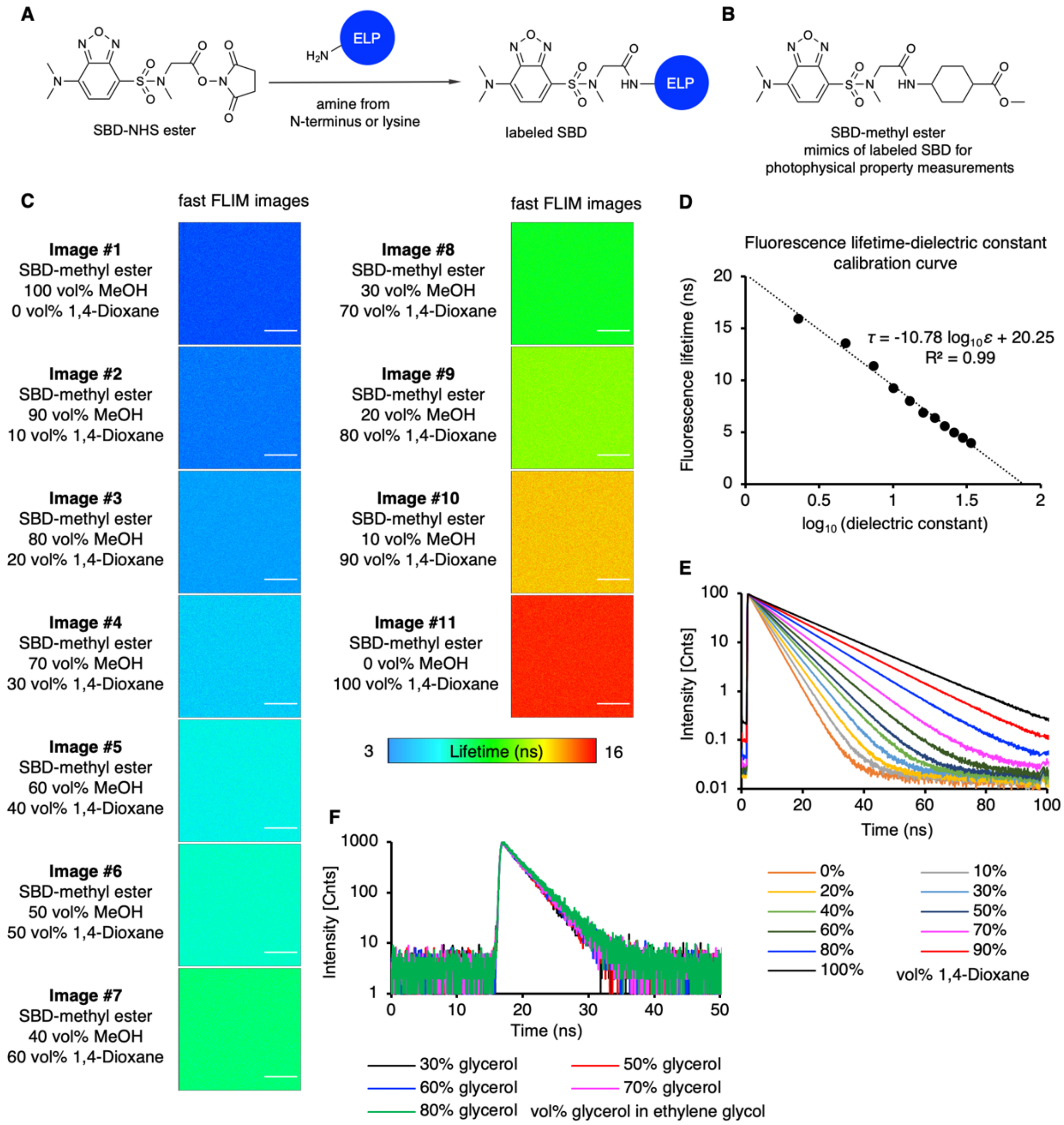


**Figure S7. Chemical structure and photophysical property of SBD. (A)** The chemical structure of free SBD-NHS probe and labeled SBD structure on the N-terminus or lysine residuals of ELP. **(B)** The chemical structure of SBD-methyl ester. SBD-methyl ester resembled the chemical structure of labeled SBD on proteins, hence was used for photophysical property measurements. **(C)** Fast FLIM images of SBD-methyl ester in solvent mixtures of methanol and 1,4-dioxane. Scale bars, 10 µm. **(D)** SBD lifetime-dielectric constant calibration curve calculated from measured SBD fluorescence lifetime in methanol and 1,4-dioxane solvent mixtures with known dielectric constant. The dielectric constant values of methanol-1,4-dioxane mixtures were calculated by the weighted average of the mixture components by assuming a simple additive effect. **(E)** Fluorescence lifetime decay curves of SBD in samples described in **C-D**. **(F)** Fluorescence lifetime decay curves of SBD in ethylene glycol-glycerol solvent mixtures demonstrated minimal fluorescence lifetime changes in comparison with SBD in methanol-1,4-dioxane mixtures. Ethylene glycol-glycerol mixtures are commonly used solvent standards bearing similar polarity but contrasting viscosities. Therefore, SBD fluorescence lifetime is insensitive to viscosity changes. The dielectric constant values of methanol-1,4-dioxane mixtures and the viscosity values of ethylene glycol-glycerol mixtures were available in **Table S3** and **S4**, respectively.

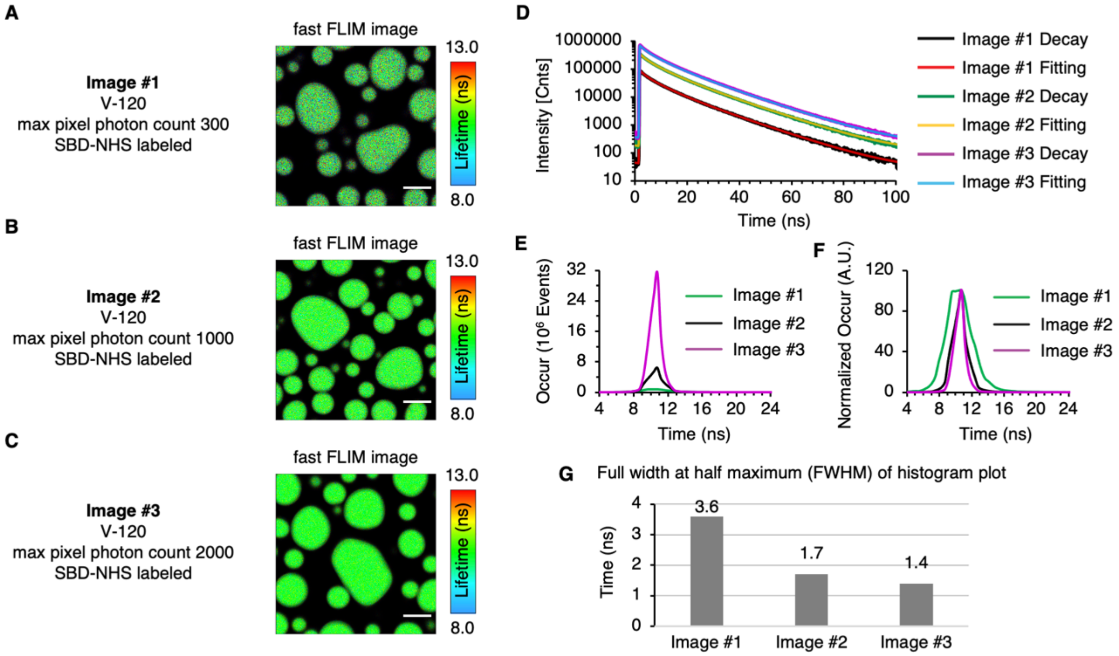


**Figure S8. FLIM images with different collected photons resulted in identical calculated fluorescence lifetime but different fluorescence lifetime distributions. (A-C)**. FLIM images of V-120 labeled with SBD-NHS with maximum photon counts 300, 1000 and 2000 per pixel, respectively. Scale bars, 5 µm. **(D)** Fluorescence lifetime decay and fitting curves from image **A-C**. **(E)** Fluorescence lifetime histogram plot from images **A-C**. **(F)** Normalized fluorescence lifetime histogram plot from image **A-C**. The peak value of each histogram curve from **A-C** was normalized to a value of 100. **(G)** Calculated FWHM plot from normalized histogram plot **F**.

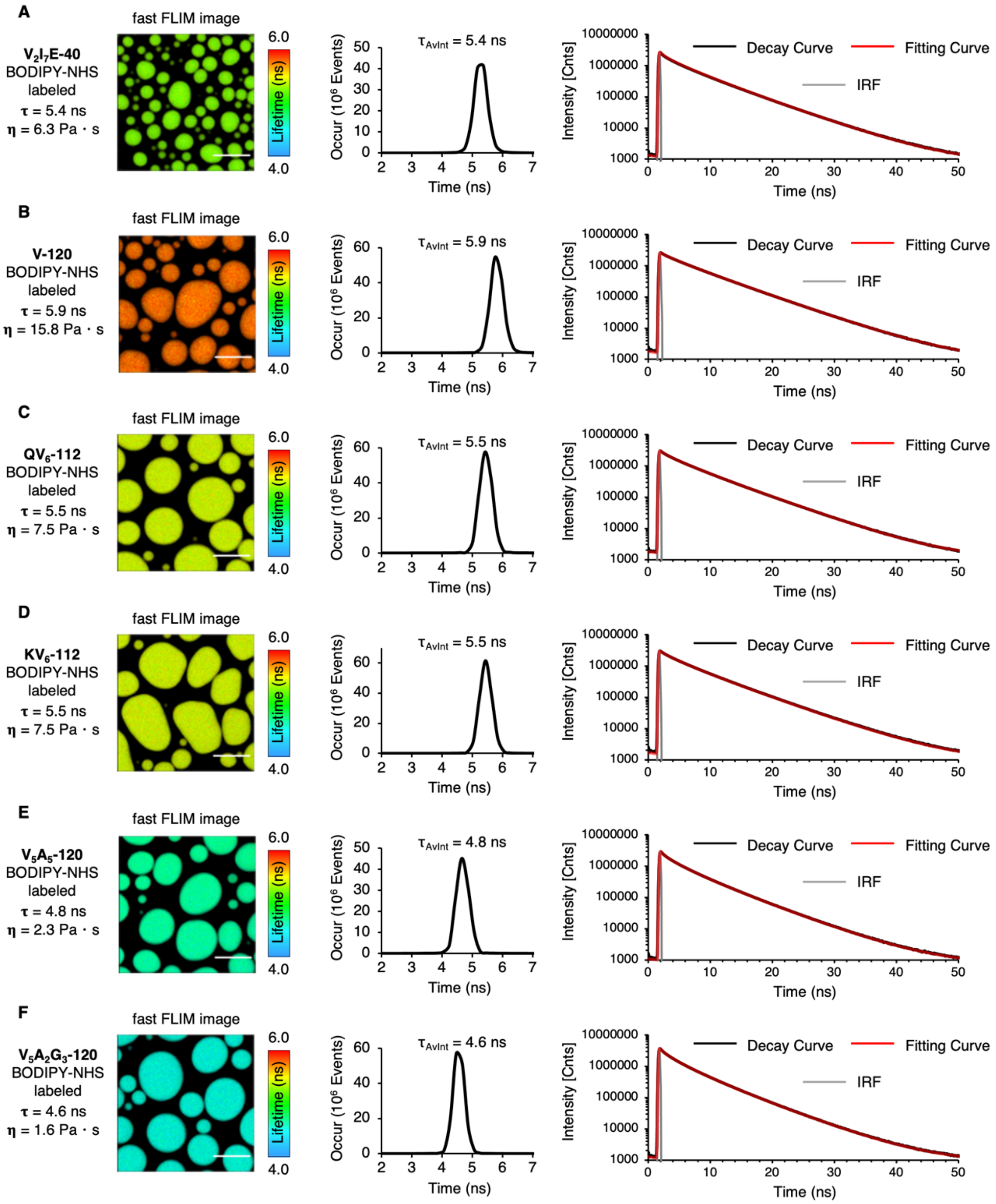


**Figure S9. Photophysical data of BODIPY-NHS labeled ELP FLIM measurements in 2 M NaCl solution. (A-F)** Fast FLIM images, fluorescence lifetime histograms and fluorescence lifetime decay curves of BODIPY-labeled ELP condensates in 2 M NaCl solution. 70 μM ELP labeled with BODIPY-NHS were mixed in phase separation buffer (50 mM HEPES, pH 7.0, 2 M NaCl) to allow FLIM imaging. Scale bars, 10 μm.

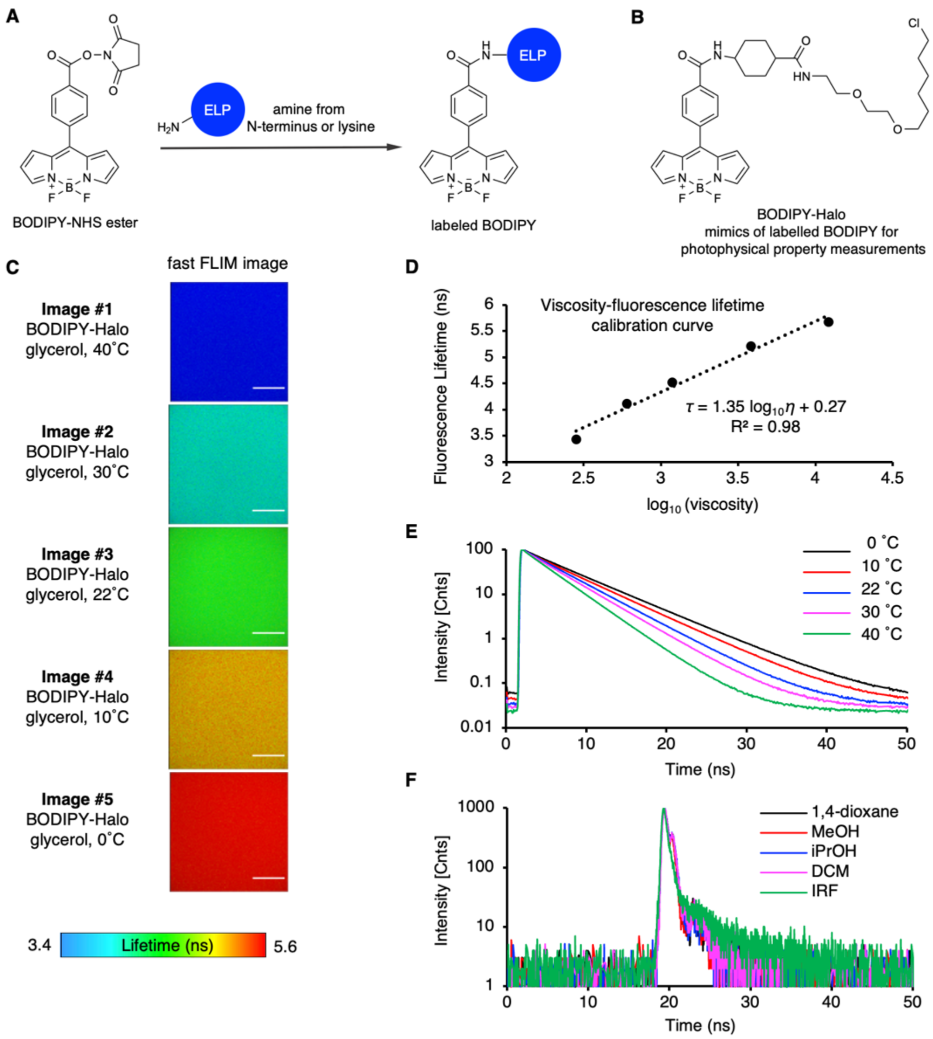


**Figure S10. Chemical structure and photophysical property of BODIPY. (A)** The structure of BODIPY-NHS probe and labeled BODIPY structure on the N-terminus or lysine residuals of ELP. **(B)** Chemical structure of BODIPY-Halo. The chromophore core part of BODIPY-Halo resembles the chemical structure of labeled BODIPY, whereas the HaloTag reactive warhead does not affect the photophysical property of BODIPY. BODIPY-Halo was used for photophysical property measurements. **(C)** Fast FLIM images of BODIPY-Halo in glycerol at different temperatures. The viscosity of glycerol is known to change dramatically in response to temperature changes, hence was used for the calibration of the BODIPY viscosity response. Scale bars, 10 μm. **(D)** BODIPY lifetime-viscosity calibration curve calculated from measured BODIPY fluorescence lifetime in glycerol at different temperatures. **(E)** Fluorescence lifetime decay curve of BODIPY-Halo in samples described in **C-D**. **(F)** Fluorescence lifetime decay curve of BODIPY-Halo in non-viscous solvents with different polarity. BODIPY displayed minimal fluorescence lifetime changes in response to different polarities. The viscosity values of glycerol at different temperatures were available in **Table S4**.

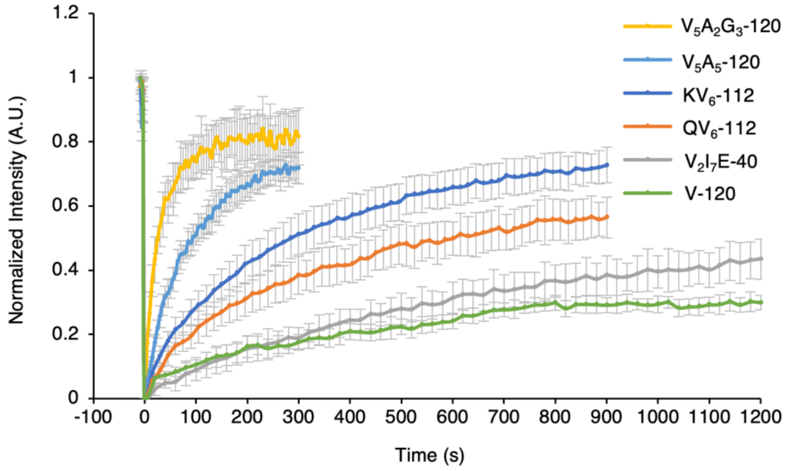


**Figure S11. Fluorescence recovery after photobleaching (FRAP) curve for ELP condensates.** 70 µM ELP labeled with fluorescein-NHS ester were mixed in phase separation buffer (50 mM HEPES, pH 7.0, 2 M NaCl) to allow FRAP measurements. Data points represent mean and SD for n = 6 for V-120; n = 7 for $V_5A_2G_3$-120, $V_5A_5$-120, $QV_6$-112 and $V_2I_7E$-40; n = 9 for $KV_6$-112.

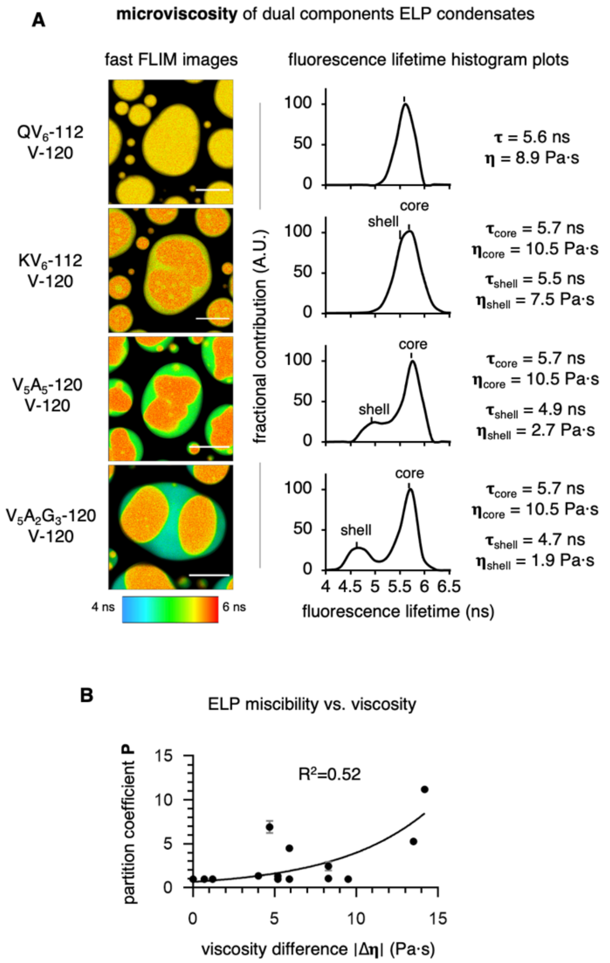


**Figure S12. The impact of microviscosity on the organization and partition of the multi-component ELP protein condensates. (A)** Pseudo-color fast FLIM images and histogram plots of dual BODIPY labeled binary ELP condensates. Scale bars, 10 µm. **(B)** Relationship between the partition coefficient of binary ELP condensates and the viscosity differences between individually formed component ELP condensates. Data were fitted to an exponential mathematical model.

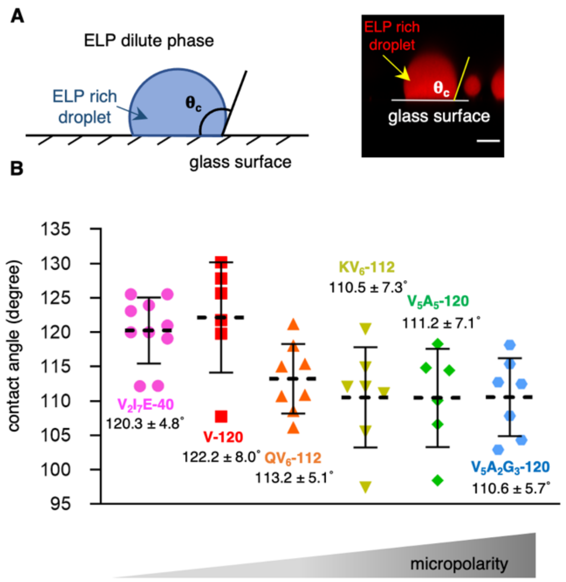


**Figure S13. Interfacial tensions of ELP condensates are reversely correlated with their micropolarity. (A)** Schematic and example of the contact angle measurements of ELP condensates on glass surfaces. Scale bar, 5 µm. **(B)** Measured contact angles of ELP condensates. 70 µM ELP labeled with AF 647 were mixed in phase separation buffer (50 mM HEPES, pH 7.0, 2 M NaCl) to allow contact angle measurements. Data points and labels represent mean and SD for n = 10 for $V_2I_7E$-40; n = 6 for V-120; n = 8 for $QV_6$-112, n = 7 for $KV_6$-112, n = 6 for $V_5A_5$-120 and n = 7 for $V_5A_2G_3$-120.

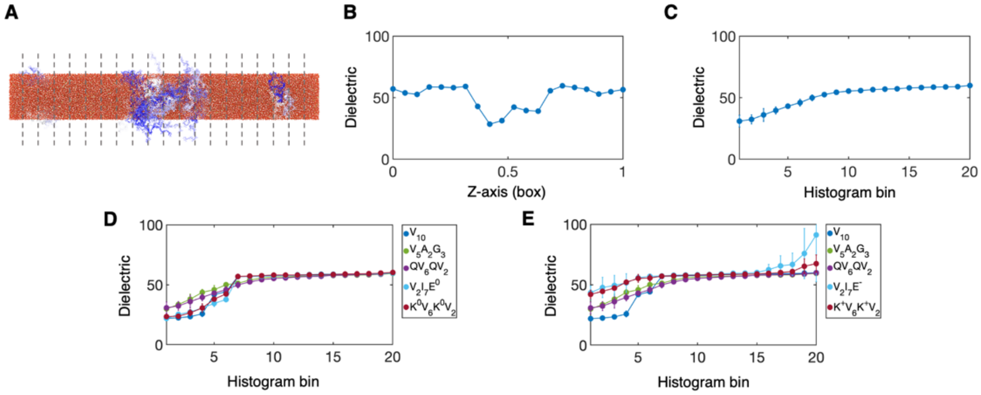


**Figure S14. Method for calculating the dielectric constant of biological condensates from molecular simulation. (A)** Final configuration of a $QV_6QV2$ all-atom simulation. The peptide is colored blue-white, water is red, $Cl^-$ is orange, and $Na^+$ is green. Grey lines represent histogram bins. **(B)** Dielectric constants are calculated for each slab of protein configurations. **(C)** These dielectric constants are sorted from lowest dielectric to highest dielectric for each simulation. **(D-E)** Three all-atom simulations are used to estimate the dielectric constant of each protein with uncharged **D** and charged **E** versions of lysine and glutamic acid. Data points represent the mean and standard deviation from three different starting configurations.

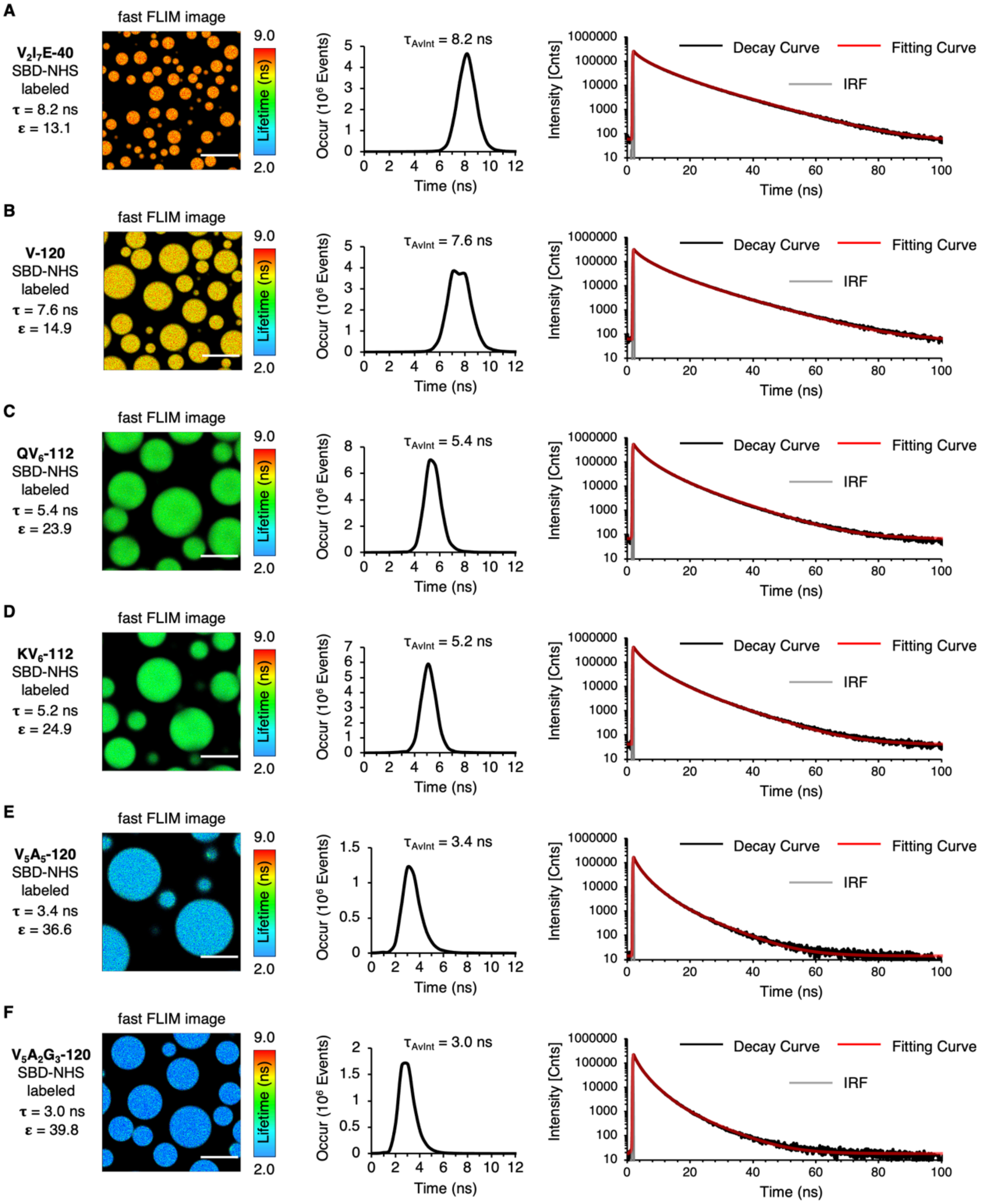


**Figure S15. Photophysical data of SBD-NHS labeled ELP FLIM measurements in 1 M NaCl solution for fitting with computational data. (A-F)** Fast FLIM images, fluorescence lifetime histograms and fluorescence lifetime decay curves of SBD-labeled ELP condensates in 1 M NaCl solution. 70 µM ELP labeled with SBD-NHS were mixed in phase separation buffer (50 mM HEPES, pH 7.0, 1 M NaCl) and incubated at 30˚C to allow FLIM imaging. Scale bars, 10 µm.

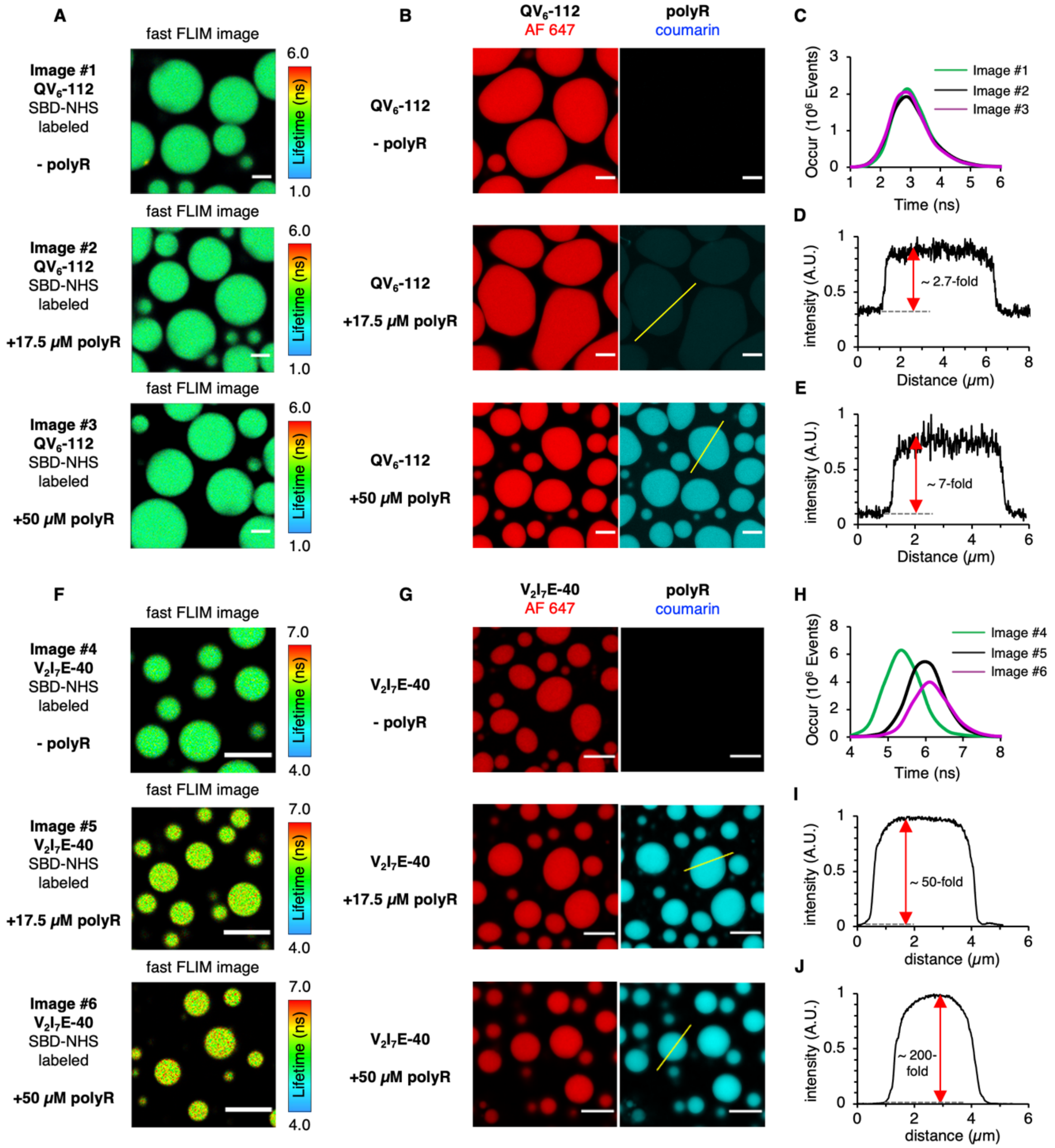


**Figure S16. Recruitment of polyR into $V_2I_7E$-40 condensate modulated its micropolarity.** PolyR is only minimally recruited to the $QV_6$-112 condensates and could not change the micropolarity of $QV_6$-112 condensates. In the case of $V_2I_7E$-40 condensates, polyR is significantly recruited to the $V_2I_7E$-40 condensates due to the strong charge-charge interaction between positive-charge polyR and negative-charge $V_2I_7E$-40. The micropolarity of $V_2I_7E$-40 condensates showed a significant decrease in the presence of polyR. **(A-B)** FLIM images of SBD-labeled $QV_6$-112 condensates (**A**) or $V_2I_7E$-40 condensates (**B**) in the absence or presence of polyR with different concentrations. **(C-D)** Dual color imaging of Alexa Fluor 647-labeled $QV_6$-112 condensates (**C**) or $V_2I_7E$-40 condensates (**D**) in the absence or presence of coumarin-labeled polyR at different concentrations. The positive charges on 17.5 μM polyR (+12 per chain) balance the negative charges on 70 μM $V_2I_7E$-40 (-3 per chain), hence the 17.5 μM polyR is the charge-neutral concentration. **(E, H)** Fluorescence lifetime histograms from images **A** and **B**, respectively. **(F-G)** Fluorescence intensity profiles along the yellow line from **C**. **(I-J)** Fluorescence intensity profiles along the yellow line from **D**. ELP phase separation condition for **A-D**: 70 µM ELP labeled with corresponding fluorophores were mixed in phase separation buffer (20 mM HEPES, pH 7.0, 150 mM NaCl) in the absence or presence of polyR at 40˚C. Scale bars, 5 µm.

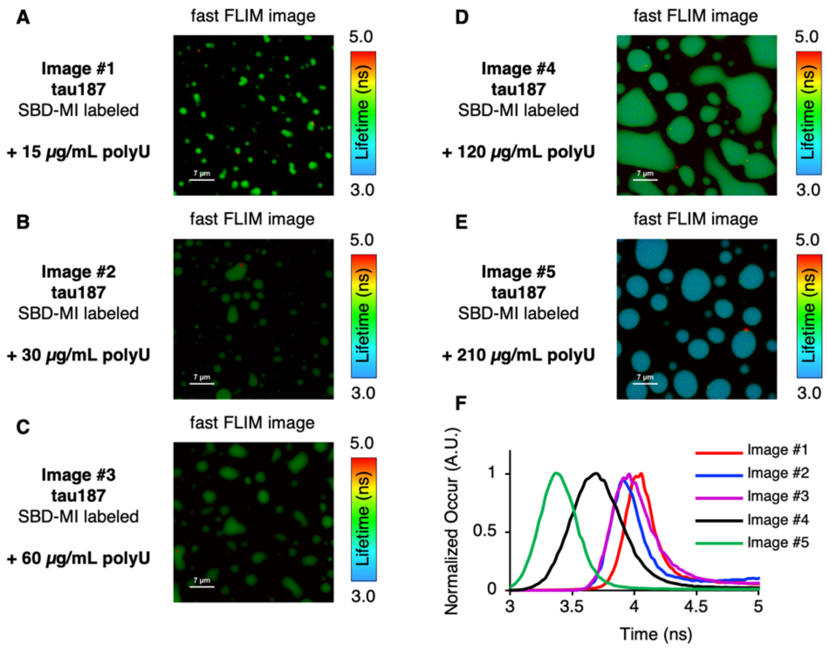


**Figure S17. The addition of RNA triggers the formation of Tau-RNA condensates through complex coacervation and dose-dependently decreases the condensates' polarity. (A-E)** Fast FLIM images of tau187 condensates in the presence of RNA. Tau187 phase separation condition: 70 µM tau187 labeled with SBD-maleimide (SBD-MI) were mixed in the phase separation buffer (20 mM HEPES, pH 7.0, 150 mM NaCl) in the presence of 15 µg/mL to 210 µg/mL polyU at 40˚C. Scale bars, 7 µm. **(F)** Normalized fluorescence lifetime histogram plot from image **A-E**.

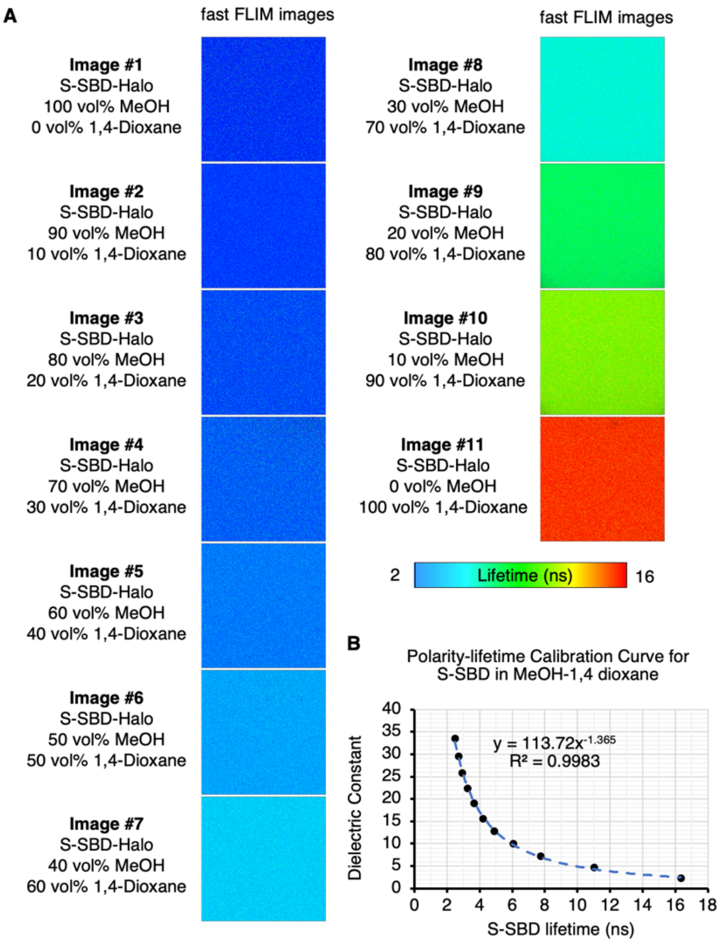


**Figure S18. Photophysical property of S-SBD-Halo. (A)** Fast FLIM images of S-SBD-Halo in solvent mixtures of methanol and 1,4-dioxane. **(B)** SBD lifetime-dielectric constant calibration curve calculated from measured SBD fluorescence lifetime in methanol and 1,4-dioxane solvent mixtures with known dielectric constant. The dielectric constant values of methanol-1,4-dioxane mixtures were calculated by the weighted average of the mixture components by assuming a simple additive effect. The dielectric constant values of methanol-1,4-dioxane mixtures was available in **Table S3**.

| | System | $\gamma$ (mN/nm) | $\varepsilon_{sim}$ | $\varepsilon_{exp}$ |
|---|---|---|---|---|
| | $V_5A_2G_3$ | -0.23 (0.54) | 30 (2) | 39.8 |
| | $QV_6QV_2$ ($QV_6$) | 0.84 (0.23) | 31 (5) | 23.9 |
| | $V_{10}$ | 1.80 (0.75) | 22 (2) | 14.9 |
| uncharged | $K^0V_6K^0V_2$ ($KV_6$) | 1.48 (0.78) | 24 (2) | 24.9 |
| | $V_2I_7E^0$ | 0.28 (0.66) | 22 (2) | 13.1 |
| charged | $K^+V_6K^+V_2$ ($KV_6$) | -0.05 (0.33) | 42 (4) | 24.9 |
| | $V_2I_7E^-$ | 0.63 (0.26) | 43 (6) | 13.1 |

**Table S1. Results for simulated surface tension and simulated dielectric constants for 10 amino acids ELP fragments in 1 M NaCl solution compared to the measured dielectric constant of ELP condensates in 1 M NaCl solution.** We note that the dielectric constants estimated from simulations of neutral K and E agree better with experimental values. Given the low polarity of the $KV_6$-112 and $V_2I_7E$ droplets, the pKa of K and E would be most likely perturbed, resulting in non-protonated K and protonated E. A similar idea has been seen in folded proteins, where interior lysine (*53*) and glutamic acid (*54*) residues are less likely to be charged. Thus, the charged $K^+$ and $E^-$ side chains were replaced with uncharged $K^0$ and $E^0$ in the simulations shown in Figure 2.

| Encoding gene | Oligo Number | Sequence from 5' end to 3' end |
|---|---|---|
| Insert with BseRI and AcuI sites | Oligo 1 | CTAGAAATAATTTTGTTTAACTTTAAGAAGGAGGAGTACATATGGGCTACTGATAATGATCTTCAG |
| | Oligo 2 | GATCCTGAAGATCATTATCAGTAGCCCATATGTACTCCTCCTTCTTAAAGTTAAACAAAATTATTT |
| V-5, fragment of V-120 | Oligo 3 | CGTGGGTGTTCCGGGCGTAGGTGTCCCAGGTGTGGGCGTACCGGGCGTTGGTGTTCCTGGTGTCGGCGTGCCGGG |
| | Oligo 4 | CGGCACGCCGACACCAGGAACACCAACGCCCGGTACGCCCACACCTGGGACACCTACGCCCGGAACACCCACGCC |
| $KV_6$, fragment of $KV_6$-112 | Oligo 5 | CGTGGGCGTACCGGGTAAAGGTGTTCCTGGCGTGGGTGTTCCGGGCGTAGGTGTCCCAGGTGTGGGCGTACCGGGCGTTGGTGTTCCTGGTGTCGGCGTGCCGGG |
| | Oligo 6 | CGGCACGCCGACACCAGGAACACCAACGCCCGGTACGCCCACACCTGGGACACCTACGCCCGGAACACCCACGCCAGGAACACCTTTACCCGGTACGCCCACGCC |
| $QV_6$, fragment of $QV_6$-112 | Oligo 7 | CGTGGGCGTACCGGGTCAGGGTGTTCCTGGCGTGGGTGTTCCGGGCGTAGGTGTCCCAGGTGTGGGCGTACCGGGCGTTGGTGTTCCTGGTGTCGGCGTGCCGGG |
| | Oligo 8 | CGGCACGCCGACACCAGGAACACCAACGCCCGGTACGCCCACACCTGGGACACCTACGCCCGGAACACCCACGCCAGGAACACCCTGACCCGGTACGCCCACGCC |
| $V_2I_7E$, first half, fragment of $V_2I_7E$-40 | Oligo 9 | CGTGGGCGTTCCGGGTATCGGTGTTCCGGGTATCGGTGTTCCGGGTATCGGTGTTCCGGGTATCGGTGTGCCGGG |
| | Oligo 10 | CGGCACACCGATACCCGGAACACCGATACCCGGAACACCGATACCCGGAACACCGATACCCGGAACGCCCACGCC |
| $V_2I_7E$, second half, fragment of $V_2I_7E$-40 | Oligo 11 | CGTGGGCGTTCCGGGTATCGGTGTTCCGGGTATCGGTGTTCCGGGTGAAGGTGTTCCGGGTATCGGTGTGCCGGG |
| | Oligo 12 | CGGCACACCGATACCCGGAACACCTTCACCCGGAACACCGATACCCGGAACACCGATACCCGGAACGCCCACGCC |
| $V_5A_5$, first half, fragment of $V_5A_5$-120 | Oligo 13 | CGTGGGTGTTCCGGGCGTGGGTGTTCCGGGTGCAGGTGTGCCGGGCGCAGGTGTTCCTGGTGTAGGTGTGCCGGG |
| | Oligo 14 | CGGCACACCTACACCAGGAACACCTGCGCCCGGCACACCTGCACCCGGAACACCCACGCCCGGAACACCCACGcc |
| $V_5A_5$, second half, fragment of $V_5A_5$-120 | Oligo 15 | TGTTGGTGTGCCGGGTGTTGGTGTACCAGGTGCAGGTGTTCCGGGTGCAGGCGTTCCGGGTGCAGGTGTGCCGGG |
| | Oligo 16 | CGGCACACCTGCACCCGGAACGCCTGCACCCGGAACACCTGCACCTGGTACACCAACACCCGGCACACCAACACC |
| $V_5A_2G_3$, first half, fragment of $V_5A_2G3$-120 | Oligo 17 | CGTGGGTGTTCCGGGCGTGGGTGTTCCGGGTGGCGGTGTGCCGGGCGCAGGTGTTCCTGGTGTAGGTGTGCCGGG |
| | Oligo 18 | CGGCACACCTACACCAGGAACACCTGCGCCCGGCACACCGCCACCCGGAACACCCACGCCCGGAACACCCACGCC |
| $V_5A_2G_3$, second half, fragment of $V_5A_2G_3$-120 | Oligo 19 | TGTTGGTGTGCCGGGTGTTGGTGTACCAGGTGGCGGTGTTCCGGGTGCAGGCGTTCCGGGTGGCGGTGTGCCGGG |
| | Oligo 20 | CGGCACACCGCCACCCGGAACGCCTGCACCCGGAACACCGCCACCTGGTACACCAACACCCGGCACACCAACACC |
| Leader sequence | Oligo 21 | TATGAGCAAAGGGCCGGG |
| | Oligo 22 | CGGCCCTTTGCTCA |
| Trailer sequence | Oligo 23 | CTGGCCGTGAGG |
| | Oligo 24 | TCACGGCCAGCC |

**Table S2. Sequences of DNA oligos used in this work.**

| MeOH Vol% | 100% | 90% | 80% | 70% | 60% | 50% | 40% | 30% | 20% | 10% | 0% |
|---|---|---|---|---|---|---|---|---|---|---|---|
| 1,4-Dioxane Vol% | 0% | 10% | 20% | 30% | 40% | 50% | 60% | 70% | 80% | 90% | 100% |
| Dielectric Constant ε | 33.6 | 29.6 | 25.9 | 22.4 | 19.1 | 15.6 | 12.9 | 10.0 | 7.3 | 4.7 | 2.3 |

**Table S3. A summary table for the dielectric constant of methanol-glycerol mixtures.**

| | | | | | | |
|---|---|---|---|---|---|---|
| Glycerol | Temperature (˚C) | 0 | 10 | 22 | 30 | 40 |
| | Viscosity (mPa·s) | 12100 | 3820 | 1180 | 600 | 280 |
| Ethylene glycol-glycerol mixture | Vol% of glycerol | 30% | 50% | 60% | 70% | 80% |
| | Viscosity (mPa·s) | 81 | 183 | 283 | 426 | 621 |

**Table S4. A summary table for the viscosity values of pure glycerol at different temperatures and the viscosity of ethylene glycol-glycerol mixtures.**

**Movie S1.**

**Sequential phase separation of AF 647-labeled V-120 and fluorescein-labeled $V_5A_2G_3$-120 from well-mixed solution as the solution temperature surpasses their $T_{ph}$.** Buffer condition, 70 μM AF 647-labeled V-120 (magenta) and 70 μM fluorescein-labeled $V_5A_2G_3$-120 (green) mixed in 20 mM HEPES, pH 7.0, 1 M NaCl. Imaging was conducted with a 1 frame/2 minutes rate. The temperature was increased from 20˚C to 40˚C with a 2˚C/min rate using a built-on heat plate on confocal microscopy.

### Reference


36. J. R. McDaniel, J. A. MacKay, F. G. Quiroz, A. Chilkoti, Recursive directional ligation by plasmid reconstruction allows rapid and seamless cloning of oligomeric genes. *Biomacromolecules* **11**, 944-952 (2010).
37. H. E. Klock, E. J. Koesema, M. W. Knuth, S. A. Lesley, Combining the polymerase incomplete primer extension method for cloning and mutagenesis with microscreening to accelerate structural genomics efforts. *Proteins Structure Function and Bioinformatics* **71**, 982-994 (2008).
38. W. Hassouneh, T. Christensen, A. Chilkoti, Elastin-like polypeptides as a purification tag for recombinant proteins. *Current Protocols in Protein Science* **61**, 6.11.1-16.11.6 (2010).
39. D. W. Peterson, H. Zhou, F. W. Dahlquist, J. Lew, A soluble oligomer of tau associated with fiber formation analyzed by NMR. *Biochemistry* **47**, 7393-7404 (2008).
40. A. Pavlova, E. R. McCarney, D. W. Peterson, F. W. Dahlquist, J. Lew, S. Han, Site-specific dynamic nuclear polarization of hydration water as a generally applicable approach to monitor protein aggregation. *Physical Chemistry Chemical Physics* **11**, 6833-6839 (2009).
41. J. Schindelin, I. Arganda-Carreras, E. Frise, V. Kaynig, M. Longair, T. Pietzsch, S. Preibisch, C. Rueden, S. Saalfeld, B. Schmid, J. Y. Tinevez, D. J. White, V. Hartenstein, K. Eliceiri, P. Tomancak, A. Cardona, Fiji: an open-source platform for biological-image analysis. *Nature Methods* **9**, 676-682 (2012).
42. A. P. Latham, B. Zhang, Maximum entropy optimized force field for intrinsically disordered proteins. *Journal of Chemical Theory and Computation* **16**, 773-781 (2019).
43. A. P. Latham, B. Zhang, Consistent force field captures homologue-resolved hp1 phase separation. *Journal of Chemical Theory and Computation* **17**, 3134-3144 (2021).
44. A. P. Latham, B. Zhang, Unifying coarse-grained force fields for folded and disordered proteins. *Current Opinion in Structural Biology* **72**, 63-70 (2022).
45. S. Rauscher, R. Pomès, The liquid structure of elastin. *Elife* **6**, e26526 (2017).
46. S. E. Reichheld, L. D. Muiznieks, F. W. Keeley, S. Sharpe, Direct observation of structure and dynamics during phase separation of an elastomeric protein. *Proceedings of the National Academy of Sciences of the U.S.A.* **114**, E4408-E4415 (2017).
47. D. A. Case, H. M. Aktulga, K. Belfon, I. Ben-Shalom, S. R. Brozell, D. S. Cerutti, T. E. Cheatham III, V. W. D. Cruzeiro, T. A. Darden, R. E. Duke, *Amber 2021*. (University of California, San Francisco, 2021).
48. G. L. Dignon, W. Zheng, Y. C. Kim, R. B. Best, J. Mittal, Sequence determinants of protein phase behavior from a coarse-grained model. *PLoS Computational Biology* **14**, e1005941 (2018).
49. Y. Zhang, S. E. Feller, B. R. Brooks, R. W. Pastor, Computer simulation of liquid/liquid interfaces. I. Theory and application to octane/water. *The Journal of Chemical Physics* **103**, 10252-10266 (1995).
50. T. A. Wassenaar, K. Pluhackova, R. A. Böckmann, S. J. Marrink, D. P. Tieleman, Going backward: a flexible geometric approach to reverse transformation from coarse grained to atomistic models. *Journal of Chemical Theory and Computation* **10**, 676-690 (2014).
51. G. A. Tribello, F. Giberti, G. C. Sosso, M. Salvalaglio, M. Parrinello, Analyzing and driving cluster formation in atomistic simulations. *Journal of Chemical Theory and Computation* **13**, 1317-1327 (2017).

52. R. J. Gowers, M. Linke, J. Barnoud, T. J. Reddy, M. N. Melo, S. L. Seyler, J. Domanski, D. L. Dotson, S. Buchoux, I. M. Kenney, in *Proceedings of the 15th python in science conference*. (SciPy Austin, TX, 2016), pp. 98-105.
53. D. G. Isom, C. A. Castañeda, B. R. Cannon, B. García-Moreno E, Large shifts in pKa values of lysine residues buried inside a protein. *Proceedings of the National Academy of Sciences of the U.S.A.* **108**, 5260-5265 (2011).
54. D. G. Isom, C. A. Castañeda, B. R. Cannon, P. D. Velu, B. García-Moreno E, Charges in the hydrophobic interior of proteins. *Proceedings of the National Academy of Sciences of the U.S.A.* **107**, 16096-16100 (2010).
